# Aphid-derived cross-kingdom RNA dynamics underpin maize resistance

**DOI:** 10.1101/2025.09.25.678037

**Authors:** Shan Jiang, Zhicheng Zhang, Chi Liu, Yeqi Zhu, Yishou Kou, Peiyu Yang, Zhichao Hu, Jun Wu, Yu Wang, Fuwei Wan, Gang Wu, Yazhou Chen

## Abstract

Aphids are major pests of maize (*Zea mays*), yet the molecular mechanisms underlying their interactions with host plants remain poorly understood. Here, we identified and functionally characterized long non-coding RNAs (lncRNAs) from the aphid-specific *Ya* gene family in the cereal-specialist aphids *Rhopalosiphum maidis* and *R. padi*. We showed that *Ya* genes formed lineage-specific clusters and were transcriptionally active across both species. Multiple *Ya* transcripts secreted into maize tissues exhibited remarkable stability compared with rapidly degraded aphid mRNAs and migrated systemically, as visualized in planta using an RNA switch-controlled RNA-triggered fluorescence system. RNA interference of *Ya* genes significantly reduced aphid fecundity, while ectopic expression of *Ya* lncRNAs in maize enhanced aphid colonization. Importantly, maize genotypes differing in aphid resistance selectively influenced the persistence of aphid mRNAs but not *Ya* lncRNAs, indicating a decoupled fate of distinct aphid RNA classes in planta. These findings establish *Ya* lncRNAs as cross-kingdom effectors critical for aphid virulence, and suggest new molecular markers for breeding aphid-resistant maize.

## Introduction

Maize (*Zea mays*) is the most widely cultivated cereal crop worldwide, and its productivity is directly tied to global food security (1–4). Among the numerous threats to maize production, aphids pose a persistent challenge because of their rapid proliferation and ability to damage plants across developmental stages. During early growth, aphid colonization reduces seedling vigor, causing chlorosis, leaf curling, and in severe cases, plant death. At reproductive stages, infestations impair pollen development and fertilization, leading to kernel abortion and yield loss (5, 6). In addition to direct feeding damage, aphid honeydew disrupts photosynthesis (6, 7), while transmission of viruses such as Maize dwarf mosaic virus (MDMV) and Barley yellow dwarf virus (BYDV) further compounds their impact (8–10). Historical outbreaks have caused yield losses of up to 50% in severely affected regions of China, and infestations are becoming increasingly frequent with the expansion of large-scale monoculture farming (11, 12).

Corn leaf aphid (*Rhopalosiphum maidis*) and the bird cherry-oat aphid (*Rhopalosiphum padi*) are the most prevalent in agricultural fields (13, 14). Their damage potential is amplified by parthenogenetic reproduction and seasonal migration between cereal crops and wild grasses (15, 16). Current management relies heavily on insecticides, but environmental concerns and regulatory restrictions underscore the need for sustainable, genetics-based alternatives (17). Breeding resistant maize varieties is the most ecologically sound strategy, yet progress has been hampered by an incomplete understanding of the molecular basis of resistance.

Early efforts to identify resistant germplasm date back more than a century, when hybrids between maize and teosinte were noted for strong aphid resistance (18). These findings suggested that domestication may have eroded innate defense traits. Subsequent screens identified resistant inbred lines (e.g., Mo17, AA8sh2, Oh545) and susceptible ones (e.g., B37, CM104, CM111) (19–21). Such studies enabled genetic mapping of resistance loci. For instance, analyses of the B73 × Mo17 recombinant inbred line (RIL) population revealed two quantitative trait loci (QTLs) on chromosomes 4 and 6 that additively enhance aphid resistance, with the chromosome 4 QTL linked to biosynthesis of 2,4-dihydroxy-7-methoxy-2H-1,4-benzoxazin-3(4H)-one (DIMBOA), a benzoxazinoid defense metabolite (22, 23). Other approaches have uncovered additional resistance loci, including a defense-related cluster in the B73 × Abe2 RIL population (24), and mutations in *LIGULELESS1* that unexpectedly confer resistance by modulating jasmonate signaling. Notably, natural variation in *LIGULELESS1* promoter activity correlates with aphid resistance, with low-expression haplotypes (Hap1, Hap2) showing enhanced defense (25).

Despite these advances, the molecular architecture of maize–aphid resistance is more complex than initially thought. For example, transposon insertions inactivate *Bx10c*, increasing the benzoxazinoid 2,4-dihydroxy-7-methoxy-1,4-benzoxazin-3-one-β-d-glucopyranose (DIMBOA-Glc) levels and enhancing resistance (22, 23). Yet, paradoxically, some susceptible lines (e.g., B73) also carry this mutation, and resistance in Mo17 can occur independently of benzoxazinoids. Additional benzoxazinoid-independent defenses have been identified, including ethylene signaling, terpenoid biosynthesis (e.g., via *Terpene synthase1*) (26, 27), and inducible flavonoid pathways (24). These layers of defense are further integrated by transcription factors such as Myeloblastosis (MYB), WRKY domain–containing (WRKY), GAI–RGA–SCR (GRAS), and NAM–ATAF–CUC (NAC) (26, 28). Still, resistance varies widely across genotypes, pointing to additional, uncharacterized mechanisms.

In parallel, aphids have evolved sophisticated strategies to overcome host defenses. Aphid feeding begins when the stylets probe plant tissues, traversing the epidermis, mesophyll, and companion cells before reaching the phloem, where prolonged sap ingestion occurs (29, 30). Throughout this process, aphids secrete saliva containing a complex mixture of proteins and RNAs that function as effectors (31–35). Some salivary proteins suppress host immunity and promote colonization (32, 34, 35). Beyond proteins, saliva also contains long noncoding RNAs (lncRNAs), such as members of the aphid-specific *Ya* gene family (33). In *Myzus persicae*, the lncRNA *Ya1* translocates into host cells, migrates systemically, and enhances aphid colonization (33).

Here, we reported the identification and functional characterization of *Ya* lncRNA family members in *R. padi and R. maidis*. We showed that multiple *Ya* transcripts were secreted into maize feeding sites and underwent systemic movement within the host. Silencing of selected *Ya* genes via RNA interference reduced aphid fecundity, whereas their overexpression in maize promoted colonization. Importantly, using both classical inbred lines (e.g., Mo17, B73) and a broader maize panel, we demonstrate that resistant genotypes attenuate aphid fecundity and alter the stability of aphid-derived RNAs. Specifically, resistance decoupled the dynamics of aphid lncRNAs and mRNAs: *Ya* transcripts remained stable across host genotypes, whereas aphid mRNAs were destabilized in resistant lines. This differential regulation highlights a previously unrecognized dimension of the maize–aphid interaction, suggesting that stable aphid lncRNAs function as effectors, while the selective destabilization of aphid mRNAs may represent a molecular basis of host resistance.

## Results

### Identification and characterization of *Ya* genes in *Rhopalosiphum* species

LncRNAs from the *Ya* gene family have been identified in *M. persicae* as virulence factors that promote aphid colonization (33). Here, we expanded the search for *Ya* genes in two *Rhopalosiphum* species, *R. padi* and *R. maidis*. Using the sequence of *MpYa1* as a query, we performed BLAST against the annotated transcripts (mRNAs and lncRNAs) of *R. padi*, *R. maidis*, and the improved annotation of *M. persicae*.

This analysis identified 21 *Ya* genes in *R. padi* (*RpYa*), 22 in *R. maidis* (*RmYa*), and 36 in *M. persicae* (*MpYa*, Fig.1A, Dataset S1-S4). Of the *MpYa* genes, 32 are located in chromosome 6 and 4 in chromosome 4 (Fig.1B). In contrast, all *Ya* genes in two *Rhopalosiphum* genomes were located in the same chromosome 2 (Fig.1B). This chromosomal distribution is consistent with known genomic rearrangements between the Macrosiphini (e.g., *M. persicae*) and Aphidini (e.g., *R. padi*, *R. maidis*) lineages (36), in which chromosomes 4 and 6 of *M. persicae* correspond to chromosome 2 of *Rhopalosiphum* species due to large-scale chromosomal fusion events during aphid speciation (Fig. S1).

**Fig. 1.**
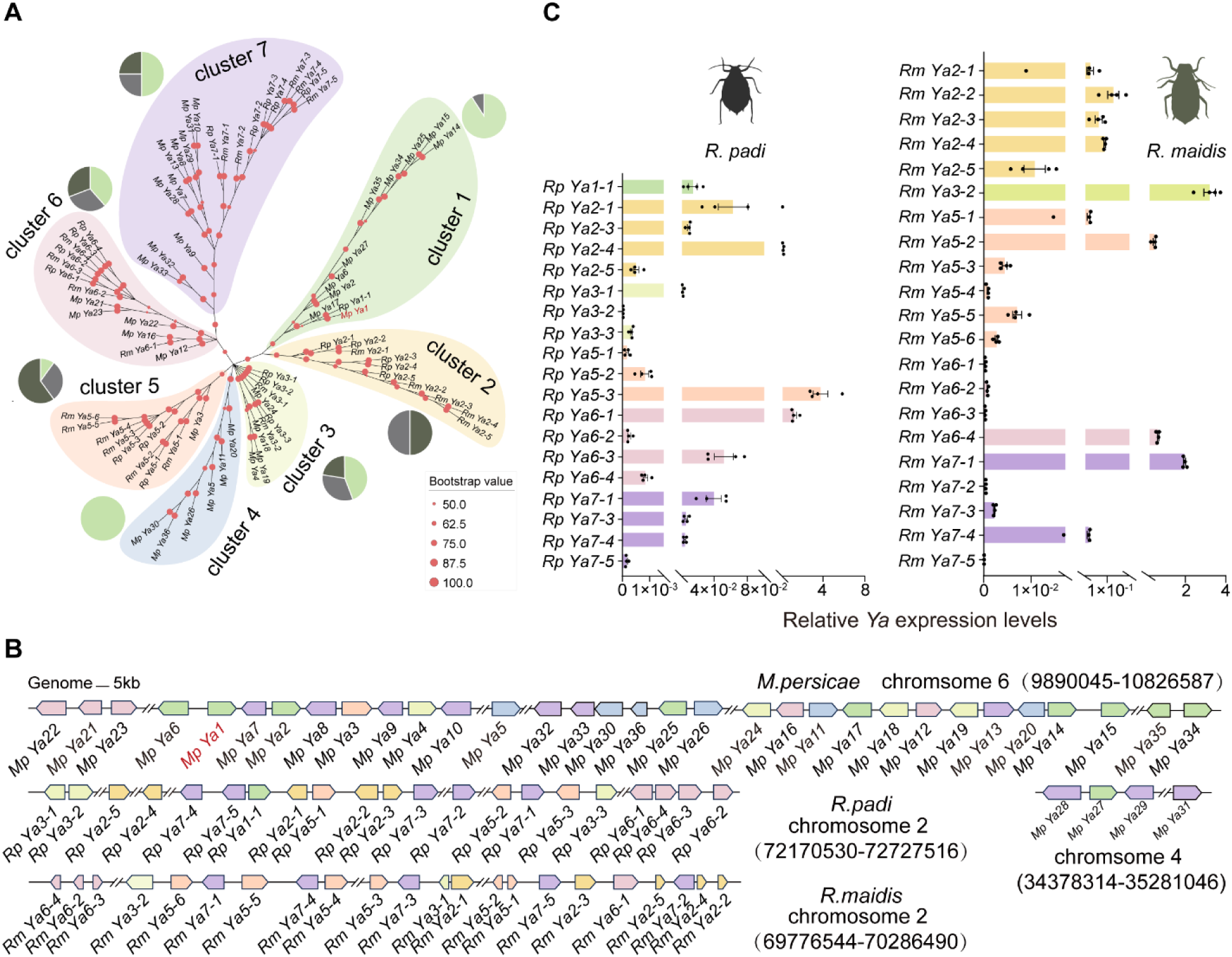
Annotation of the *Ya* gene family in the genomes of *M. persicae*, *R. padi*, and *R. maidis*. (A) Bayesian phylogeny of *Ya* genes from *M. persicae* (n = 36), *R. padi* (n = 21) and *R. maidis* (n = 22). The pie charts illustrate the proportional distribution of *Ya* genes within each cluster, with light green representing *M. persicae*, gray for *R. padi*, and dark green for *R. maidis*. (B) *Ya* genes are tandemly duplicated genes locate closely on the chromosomes of each aphid species. Arrows indicated the transcription orientation, colors indicated phylogenetic clusters shown in Fig. 1A. (C) Expression levels of *Ya* genes in three aphid species via RT-PCR. Colors indicated phylogenetic clusters shown in Fig. 1A. A total of four replicates were analyzed. Each replicate consisted of 50 aphids feeding on a 4-leaf-stage B73 maize plants. Expression levels of *RpYa* and *RmYa* genes were related to the house-keeping genes *R. padi Ribosomal protein L7* (*RpRpL7*) and *R. maidis Elongation factor 1-α* (*EF1α*), respectively.

Phylogenetic analysis grouped the *Ya* genes into eight distinct clusters (Fig.1A). Cluster 2 and Cluster 4 were species-specific, containing only *Ya* genes from either *R. padi* and *R. maidis*, or from *M. persicae*, respectively. In contrast, Cluster 1 and Cluster 6 were dominated by *Ya* genes from either *M. persicae* or the Aphidini species. Clusters 3, 7, and 8 included *Ya* genes from all three species, with subclades in Clusters 7 and 8 further separating into *M. persicae*- and *Rhopalosiphum*-specific lineages, suggesting both conserved and lineage-specific expansion patterns. Gene expression analysis confirmed that the *Ya* genes identified in *R. padi* and *R. maidis* are transcriptionally active (Fig.1C), although their expression levels varied markedly across species and phylogenetic clusters.

### Cross-kingdom translocation of *Ya* lncRNAs from *Rhopalosiphum* species into maize plants

*MpYa1*, *MpYa2*, *and MpYa17* were previously found to be translocated and systematically migrate in various plants (33). Notably, *MpYa1* was also found to be translocated into maize plants via *M. persicae* feeding (33). These three *Ya* genes were in Cluster 1, with the other seven *Ya* genes from *M. persicae* and only from *R. padi* (*RpYa1-1*) (Fig. 1A). Intriguingly, *RpYa1-1* belonged to a subclade with *MpYa1* and *MpYa17*, suggesting *RpYa1-1* was likely to be a cross-kingdom RNA.

To test whether *RpYa1-1* was translocated into maize plants, we set up the feeding experiments. 50 *R. padi* were caged on maize leaves for 48 hours (Fig. 2A). The caged parts were the feeding sites (FS), and the uncaged tissues above and below were the up- feeding sites (UFS) and down-feeding sites (DFS), respectively. The controls were the counterparts on the maize plants without aphid treatment. Leaves were carefully washed and processed for RNA extractions. PCR on FS, UPS, and DFS revealed that *RpYa1-1* was detected in those sites (Fig. 2B), confirming that *RpYa1-1* was a cross-kingdom RNA. Sequences of PCR products obtained from FS, UFS, and DFS confirmed the presence of *RpYa1-1* translocated into maize plants (Fig. S2A).

**Fig. 2.**
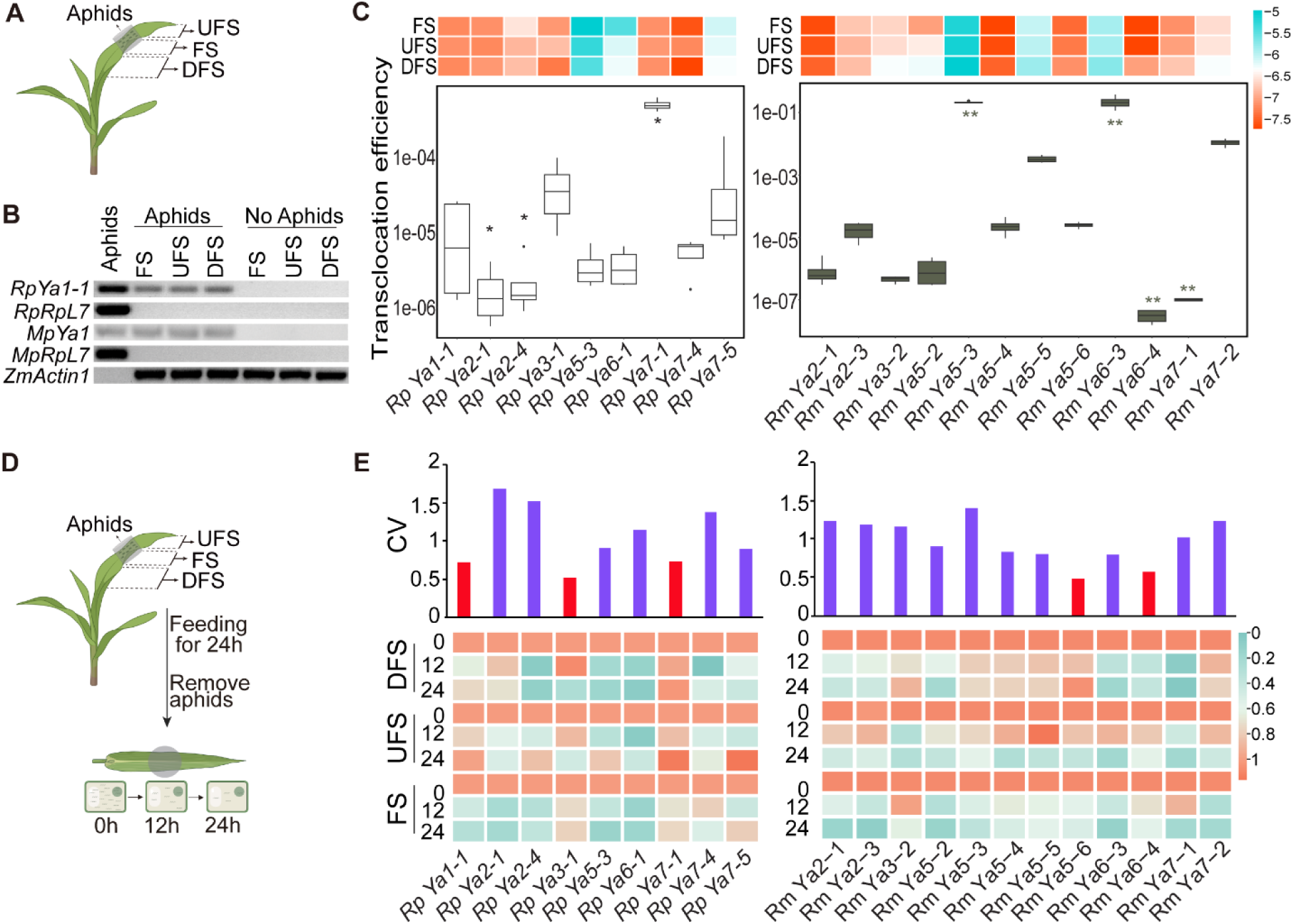
*Ya* lncRNAs were selectively translocated into maize plants and few exhibited high stability. (A) Experimental setup for aphid feeding. 50 aphids were confined to the leaves of 4-leaf-stage maize plants for 48 h. The caged region was designated as the feeding site (FS), while adjacent regions were defined as UFS and DFS. The counterparts on the uninfested maize plants with empty caged on the FS were harvested as controls. (B) Detection of *Ya* transcripts in maize tissues. *R. padi RpYa1-1* and *M. persicae MpYa1* were translocated into maize and systematically migrated to distal tissues. RT-PCR using specific primers confirmed the presence of *RpYa1-1* and *MpYa1* in FS, UFS, and DFS. Housekeeping genes (*MpRpL7*, *RpRpL7*, and *ZmActin1*) were used as controls. (C) Relative abundance and translocation efficiency of *Ya* transcripts. Cross-kingdom *RpYa* and *RmYa* abundance in maize plants were quantified by RT-PCR and normalized to *ZmActin1*(boxplots of relative abundance are shown in Fig. S6). Subsequently, the transcript abundance at each time point and in each tissue site was normalized relative to the abundance at the 0 hour time point and visualized as a heatmap. Translocation efficiency was calculated as the ratio of transcript abundance in FS to that in aphids. Aphid transcript levels were normalized to *RpRpL7* (*R. padi*) or *RmEF1α* (*R. maidis*). Statistical differences in translocation efficiency among genes were assessed with the Kruskal–Wallis test followed by Dunn’s post hoc test (Bonferroni correction). Significance levels are indicated as \**p* < 0.05 and \*\**p* < 0.01. Abundance data for aphids and FS are provided in Fig. 5. (D) Experimental setup for stability assays. Using the same feeding system as in panel (A), aphids were removed after 48 h. Maize plants samples were collected at 0, 12, and 24 h after aphid removal. (E) Stability and spatiotemporal dynamics of *Ya* transcripts. Transcript abundance across time points and tissue sites was quantified as in panel (C). Stability was assessed by calculating the coefficient of variation (CV) across biological replicates. Low CV values indicate stable expression, shown by red bars. All CV values are provided in Table S2.

Since *RpYa1-1* was the only *Ya* from *Rhopalosiphum* species clustered with *MpYa1*, it indicated that those two species may have different translocation profiles of *Ya* RNAs compared to *M. persicae*. Therefore, we investigate whether the *Ya* homologs in two aphid species were translocated into the maize plants. Cage experiments that involved two species were set up on maize leaves (Fig. 2A), as described above. With the gene-specific primers (Table S1), we found 9 additional *RpYa* and 12 *RmYa* were detected in the FS, UFS, and DFS (Fig. S3). Migration of these *Ya* lncRNAs was further confirmed by the sequences of PCR products obtained by amplification using cDNA of FS, UFS, and DFS (Fig. S4). Interestingly, compared to the expression levels in aphids and translocated *Ya* (Fig. 2C, Fig. S5), the cross-kingdom ability was unlikely to be related to their abundance in aphids.

### Stability of aphid-derived *Ya* RNAs in maize plants

As foreign RNA molecules, aphid-derived *Ya* RNAs are expected to encounter plant defenses that degrade exogenous RNAs, such as antiviral RNA mechanisms (37). To assess the stability of aphid *Ya* RNAs in maize plants, we conducted a time-course experiment in which *R. padi* and *R. maidis* were allowed to feed on maize leaves for 48 hours, after which the aphids were removed. Leaf samples from FS, as well as UFS and DFS, were collected at 0、12 and 24 hour intervals post-removal (Fig. 2D).

The presence and abundance of *Ya* transcripts in FS, UFS, and DFS were investigated by RT-PCR. Most *Ya* transcripts from both aphid species showed a time-dependent decline in abundance at the FS (Fig. 2E, Fig. S6A and B), indicating progressive degradation. Notably, the serval *Ya* remained detectable at relatively high levels over time (Fig. 2E; Fig. S6A and B), suggesting higher stability compared to other *Ya* RNAs. These differences became more obvious in the UFS, where most *Ya* RNAs showed sharp declines (Fig. S6A and B). Coefficient of variation analysis on Y*a* abundance across three time points and three sites revealed that *RpYa3-1*, *RpYa1-1*, *RpYa7-1* in *R. padi* and *RmYa5-6* in *R. maidis* were relatively stable in the FS compared to other *Ya* lncRNAs (Fig. 2E, Table S2).

Together, these results indicate that while most aphid-derived *Ya* RNAs are subject to degradation in maize tissue, a subset exhibits relative persistence, potentially enabling functional interactions within the host.

### Visualization of *Ya* localization in planta

Considering both translocation efficiency and stability, we chose *RpYa7-1* and *RmYa5-6* for further investigation. We examined the in planta migration of *RpYa7-1* and *RmYa5-6* using an RNA imaging approach, the RNA switch–controlled RNA-triggered fluorescence (RNA switch–RTF) system, which enables real-time visualization of RNA dynamics in living plants (38). In this system, the RNA switches with probes targeting either *RpYa7-1* or *RmYa5-6* can toggle between active and inactive states depending on the presence of the target RNA, thereby controlling GFP accumulation or degradation (Fig. 3A). To design a specific probe, we first determined the full-length *RpYa7-1* (469 nt) and *RmYa5-6* (394 nt) sequences using 3′ RACE (Fig. S7A). Based on this sequence, we designed suitable RNA switches with the probes for both *Ya* (Fig. S7B).

**Fig. 3.**
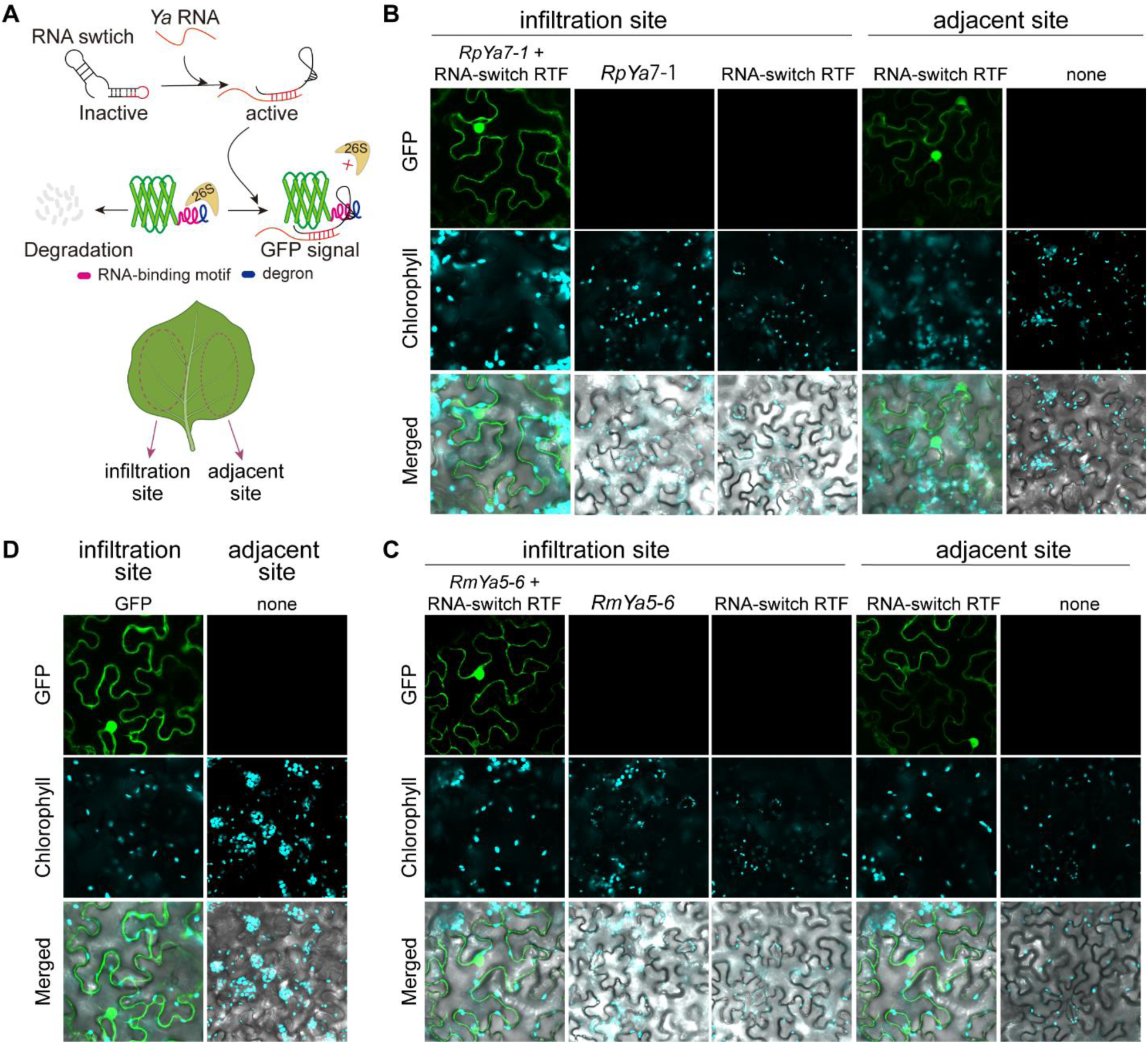
Visualization of *RpYa7-1* and *RmYa5-6* localization and movement. (A) Schematic of the RNA-switch–RTF reporter system. The red sequence in RNA-switch represents the probes for *RpYa7-1* or *RmYa5-6*. In the absence of target RNA, the RNA-switch remains inactive due to base pairing between the linkers and their complementary sequences in the probe, leading to GFP degradation via 26S proteasome. Upon target RNA binding, the probe hybridizes with the target, triggering the switch to an unfolding active state, recruiting in GFP accumulation at the RNA site (For details, please refer to the Materials and Methods section). (B) Schematic of infiltration injection site in *N. benthamiana* leaves. (C) Visualization of *RpYa7-1* or *RmYa5-6* localization and movement in *N. benthamiana* leaves using the RNA-switch–RTF system. The infiltration site was injected with either a mixture of the *RpYa7-1* or *RmYa5-6* expression construct and the RNA-switch–RTF construct, or with the RNA-switch–RTF construct alone, while the adjacent site was treated with the RNA-switch–RTF construct accordingly. (D) GFP expressed alone was unable to move in *N. benthamiana* leaves. None indicated no construct infiltrated.

Agroinfiltration of *N. benthamiana* leaves with AtUbq10::RpYa7-1 or AtUbq10::RmYa5-6 together with the corresponding RNA switch–RTF construct produced GFP signals in both the nucleus and cytoplasm (Fig. 3B and C), confirming subcellular localization of *RpYa7-1* and *RmYa5-6*. No GFP fluorescence was observed when leaves were infiltrated with either *Ya* or the RNA switch–RTF system alone (Fig. 3B and C). Remarkably, GFP signals were also detected in regions adjacent to the infiltration sites, where the RNA switch–RTF system had been introduced but not the AtUbq10::RpYa7-1 or AtUbq10::RmYa5-6 plasmid (Fig. 3B and C), demonstrating systemic movement of *RpYa7-1* and *RmYa5-6*. By contrast, infiltration with 35S::GFP produced strong local fluorescence but no signals in neighboring tissues (Fig. 3D).

Together, these results demonstrate that *RpYa7-1* and *RmYa5-6* are not only localized within plant cells but also migrate systemically from the initial delivery sites to adjacent and distal tissues.

### Translocated aphid lncRNAs are important for aphid colonization

To investigate the role of aphid *Ya* lncRNAs in host colonization, we performed gene expression knockdown experiments by dsRNA feeding, targeting *RpYa7-1* in *R. padi* and *RmYa5-6* in *R. maidis*, respectively (Fig. 4A and E). Knockdown of these lncRNAs significantly reduced aphid fitness on maize, as evidenced by decreased nymph production (Fig. 4B and F) and lower honeydew secretion (Fig. 4C and G). In addition, RNAi-treated aphids showed markedly reduced survival rates compared to control groups (Fig. 4D and H). These findings suggest that *RpYa7-1* and *RmYa5-6* are critical for the successful colonization of maize by *R. padi* and *R. maidis*, respectively.

**Fig. 4.**
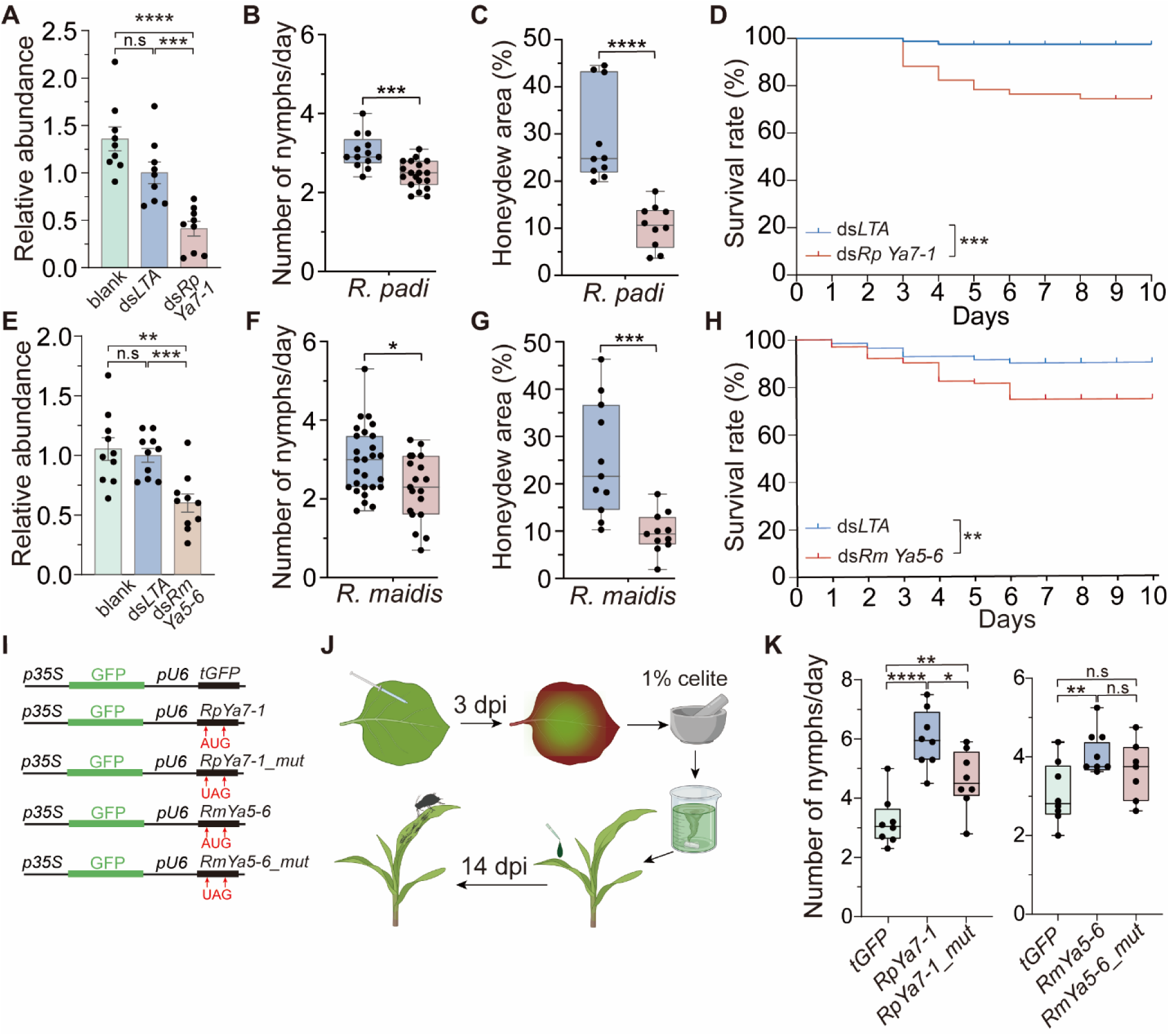
*RpYa7-1* and *RmYa5-6* promote aphid colonization on maize. (A, E) Knock-down the expression of *RpYa7-1* in *R. padi* and *RmYa5-6* in *R. maidis* by dsRNA feeding. Transcript levels were normalized to *RpRpL7* and *RmEF1α* (mean ± SD, n = 9–10). (B, F) RNAi-aphids produced fewer nymph. Each dot represents one aphid (n = 12–27; one aphids per maize plant). (C, G) RNAi-aphids secreted less honeydew. Honeydew was collected for 48 h on filter paper, stained with aniline blue, and quantified by ImageJ (n = 10–12; five aphids per maize plant). (D, H) Knockdown of *Ya* decreased aphid survival. Survival was monitored daily and compared between treatments after RNAi over 10 days. (I) Schematic of the FoMV plasmid designed for virus-mediated lncRNA overexpression in maize. A U6 promoter-driven module containing the full-length *RmYa5-6* lncRNA was inserted downstream of the EGFP reporter gene. In addtion, mutated sequences (AUG-to-UAG) were aslo generated, *RmYa5-6-mut*, and inserted downstream of the EGFP. Control constructs contained a truncated GFP (tGFP) is a non-coding EGFP fragment (76–495 nt). (J) Experimental workflow for FoMV-mediated gene expression in maize. *N. benthamiana* was agroinfiltrated to produce viral particles, which were mechanically inoculated onto maize leaves. EGFP fluorescence at 14 dpi confirmed systemic infection (Fig. S8C and D). (K) Overexpression of *RpYa7-1 and RmYa5-6* enhanced aphid fecundity. Each dot represents the total offspring from a single aphid placed on a systemically infected maize plant (n = 7–8 plants; one aphid per plant). Data in B, C, F, G, and K were analyzed by an unpaired two-tailed Student’s t-test. Survival curves in D and H were analyzed by the Kaplan-Meier method and compared using the log-rank (Mantel-Cox) test. Significance levels: ns, not significant; \**p* < 0.05; \*\**p* < 0.01; \*\*\**p* < 0.001; \*\*\*\**p* < 0.0001. The value of n indicates the sample size for each experiment.

To further test whether these lncRNAs also function in *planta* to promote aphid colonization, we intended to ectopically express *RpYa7-1* and *RmYa5-6* in maize plants. Bioinformatic analysis identified a short open reading frame (ORF) in each lncRNA. To eliminate the potential confounding effects of peptide translation, we generated mutated versions of the lncRNAs in which all AUG start codons were replaced with UAG stop codons (Fig. 4I). As a control, we included a truncated version of GFP (tGFP), composed of a 420-nt GFP fragment lacking a start codon (Fig. 4I).

For overexpression, we employed a virus-mediated delivery system using a modified PV101 vector (39). In this system, GFP protein expression was driven by the CaMV 35S promoter, while lncRNA sequences (*RpYa7-1*, *RpYa7-1-mut*, *RmYa5-6*, *RmYa5-6-mut*, and *tGFP*) were driven by the U6 promoter. The recombinant PV101 plasmids were first agroinfiltrated into *N. benthamiana* leaves (Fig. 4J). Three days post-infiltration (dpi), infected leaves were ground, and the sap was rubbed onto maize leaves at the 3-leaf stage. At 14 dpi, systemic GFP fluorescence was observed throughout the maize plants, including in the roots, confirming successful viral spread (Fig. 4J, Fig. S8). RT-PCR confirmed robust expression of both native and mutant *Ya* lncRNAs in corresponding overexpression plants, whereas no *Ya* transcripts were detected in the *tGFP* control plants (Fig. S9). Importantly, coat protein RNA levels were comparable across all treatment groups, indicating similar infection efficiency.

When aphids fed on the overexpression plants, *R. padi* on RpYa7-1 and RpYa7-1-mut lines, and *R. maidis* on RmYa5-6 and RmYa5-6-mut lines, produced significantly more offspring (Fig. 4K) compared with aphids feeding on *tGFP* control plants. These results indicate that both the native and AUG-mutated forms of *Ya* lncRNAs can enhance aphid performance, supporting the conclusion that *RpYa7-1* and *RmYa5-6* act as transcripts promoting aphid colonization, independent of their coding potential.

### Aphid mRNAs migrated systematically in the maize plants and were degraded rapidly

Translocation of *Ya* RNAs to maize plants is likely through aphid saliva, which also contains other RNAs, such as aphid mRNAs. To investigate the presence of aphid mRNAs in the FS, we conducted RNA-seq analysis on FS samples fed *R. padi* and *R. maidis*, respectively. The controls were the maize leaves that were untreated with aphids (Fig. 5A). Reads were mapped to a merged genome file containing the genomes of *R. padi* or *R. maidis* together with the maize B73 reference genome. The vast majority of mapped reads (>99.9%) derived from maize RNAs, and average 0.01% reads in the FS fed by *R. padi* and 0.09% of reads in the FS fed by *R. maidis* derived from aphids (Fig. 5B). From those mapped reads, 2424 *R. padi* RNAs and 5774 *R. maidis* RNAs were detected. Of which, 8 *RpYa* and 10 *RmYa* RNAs were present in at least one FS replicate (Datasets S5 and S6). The translocation of three *RpYa* (*RpYa5-3*, *RpYa6-1*, *RpYa7-1*) and five *RmYa* (*RmYa3-2*, *RmYa5-5*, *RmYa5-6*, *RmYa6-3*, *RmYa7-*1) in the maize leaves was consistent with the results above (Fig. 2).

**Fig. 5.**
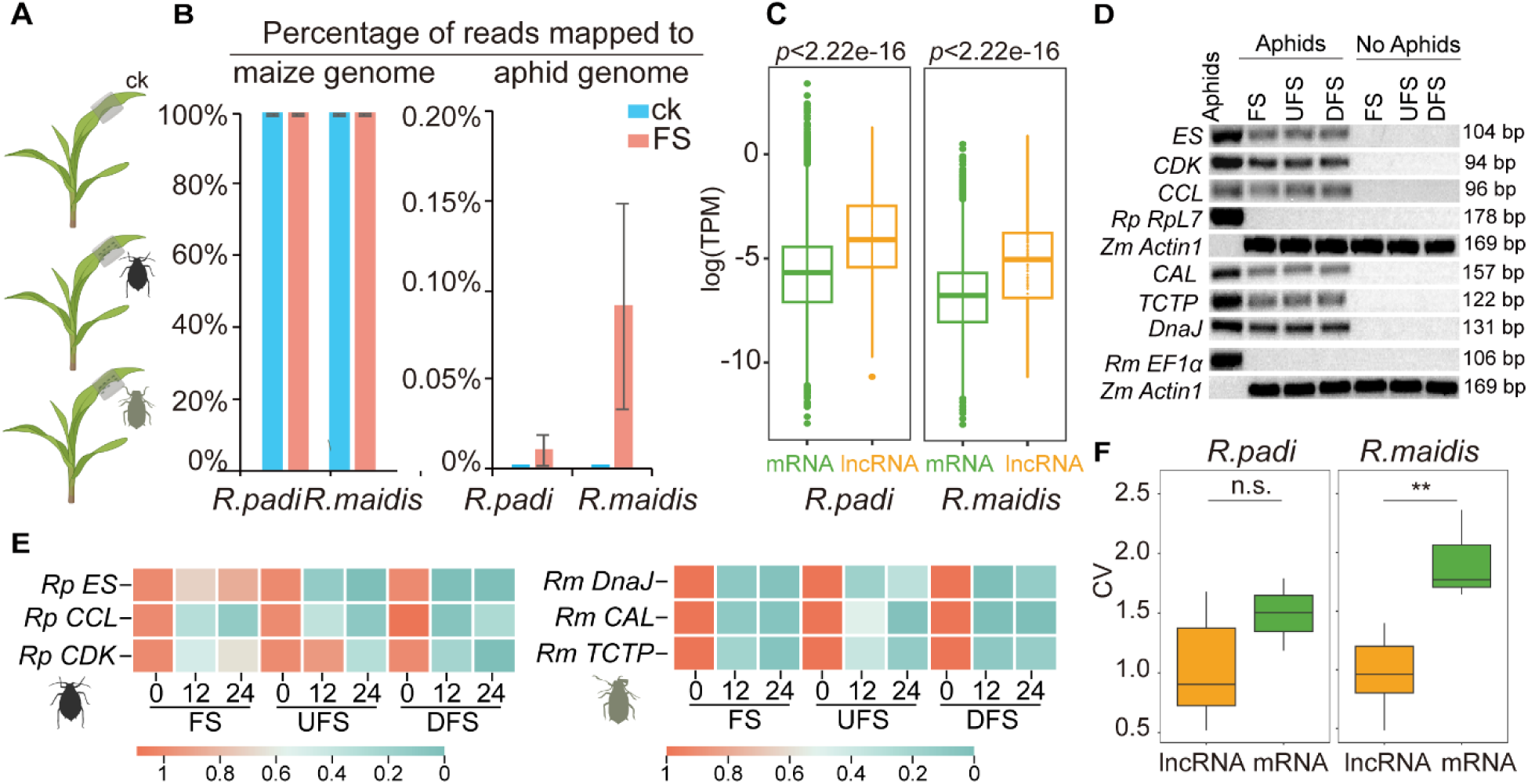
Aphid mRNAs are secreted into maize but are less stable than *Ya* lncRNAs. (A) Experimental design for transcriptome analysis of FS. (B) Mapping of transcriptome reads to maize and aphid genomes. Bars showing the percentage of RNA-seq reads aligned to the maize genome versus the aphid genome. (C) Aphid-secreted lncRNAs are more abundant than mRNAs at the FS. Box and scatter plots showing log(TPM) values of mRNAs (green) and lncRNAs (orange) fro*m R. padi* (left) and *R. maidis* (right). (D) Detection of mRNAs in maize tissues. *R. padi* (*RpES*、*RpCDK*、*RpCCL*) *R.maidis* (*RmCAL*、*RmTCTP*、*RmDnaJ*) mRNAs were translocated into maize and systematically migrated to distal tissues. RT-PCR using specific primers confirmed the presence of these mRNAs in FS, UFS, and DFS. Housekeeping genes (*RpRpL7*, *RmEF1α* and *ZmActin1*) were used as controls. (E) Stability and spatiotemporal dynamics of cross-kingdom mRNAs. These mRNAs abundance was quantified by RT-PCR and analyzed as in Fig. 2E. Relative abundance is shown as bar plots in Fig. 10. (F) Stability comparison of mRNAs and lncRNAs. Box plots showing the CV for lncRNAs (orange) and mRNAs (green) from *R. padi* (left) and *R. maidis* (right). n.s. indicate *p* >0.05; ** *p* < 0.01, unpaired two-tailed t-test), determined by the Wilcoxon rank-sum test.

In addition to *Ya* RNAs, both aphid species secrete large numbers of their RNAs into the FS on the maize leaves. Overall TPM values of aphid mRNAs detected in the FS were significantly lower than those of aphid lncRNAs (Fig. 5C). We selected three representative mRNAs from each species, *ES*, *CDK, CCL* from *R. padi*, and *CAL*, *TCTP*, and *DnaJ* from *R. maidis* (Datasets S5 and S6), then tested their presence in the FS on maize plants (Fig. 5D). We found that these mRNAs were detected in the FS, UFS, as well as DFS, suggesting the systematic migration of aphid mRNAs in the maize plants. We also found that the abundances of these mRNAs were dramatically decreased over time in the FS after aphids were removed. These trends were more obvious in the UFS and DFS samples (Fig. 5E).

### Decoupled stability of aphid lncRNAs and mRNAs in maize reveals the molecular basis of plant resistance

To evaluate the impact of host plant resistance on aphid-delivered cross-kingdom RNAs, we first classified maize genotypes based on aphid reproduction. As expected, the resistant line Mo17, previously characterized as resistant to *R. padi* and *R. maidis*, supported significantly lower aphid fecundity compared with the susceptible reference line B73 (19–21) (Fig. 6A and C). Screening an additional 20 lines from the CUBIC (Complete-diallel design plus Unbalanced Breeding-like Inter-Cross) population (40) (Fig. S11) identified four additional resistant lines (E28, Ji853, Zi330, H21) and one highly susceptible line (Nx110) (Fig. 6B and D).

**Fig. 6.**
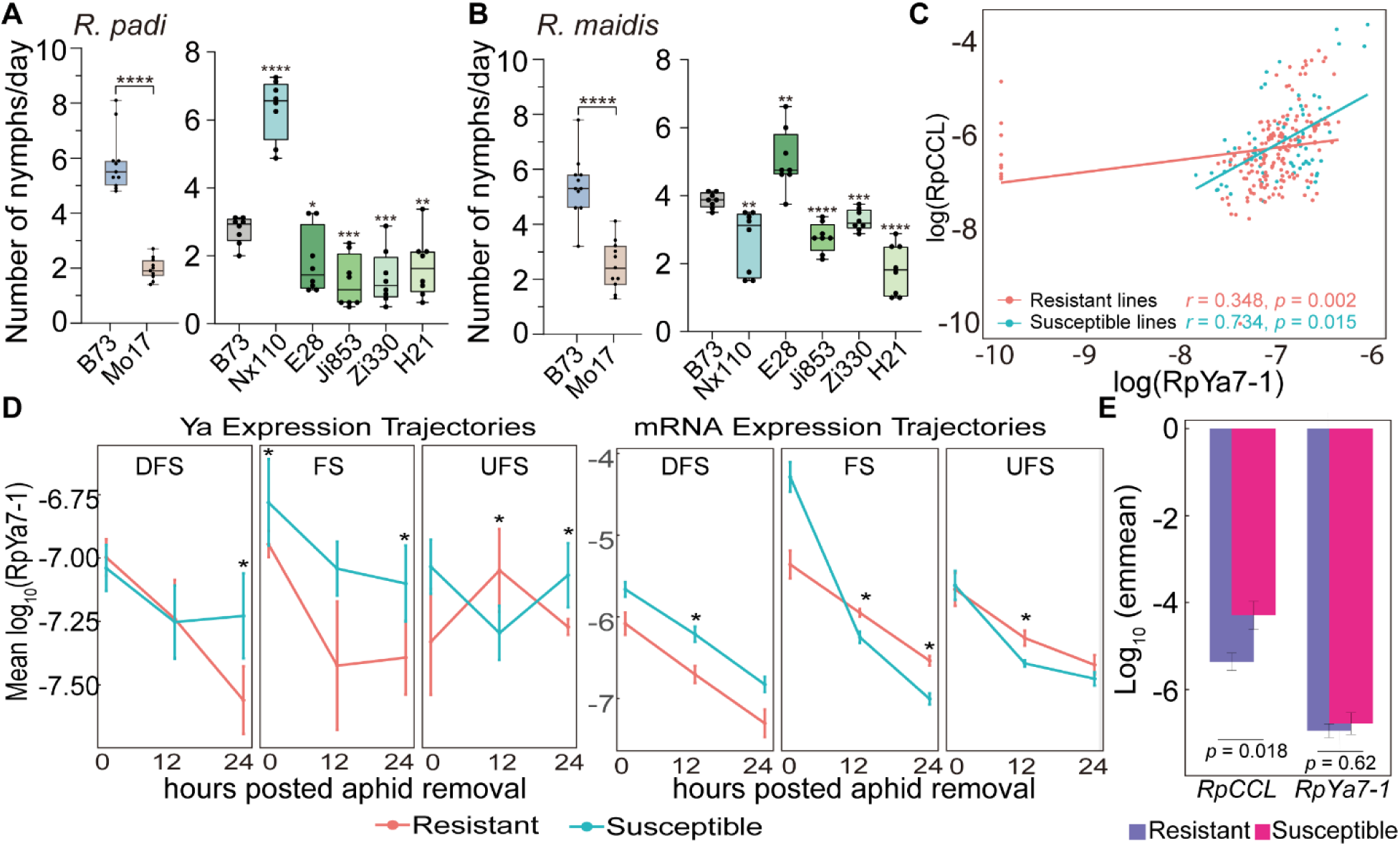
Maize resistance decouples the stability of aphid-delivered lncRNAs and mRNAs. (A, B) Fecundity of *R. padi* (A) and *R. maidis* (B) on different maize varieties (Nx110, E28, Ji853, Zi330, H21 from the CUBIC population). Each dot represents the daily offspring number from a single aphid (n = 8–11 plants per variety). Significance was determined by unpaired two-tailed t-tests against B73: ns, not significant; *p* < 0.05; *p* < 0.01; *p* < 0.001; *p* < 0.0001 (C) Correlation between aphid-derived *RpYa7-1* (lncRNA) and *RpCCL* (mRNA) abundance. Correlation was strong in susceptible maize lines (r = 0.734) but weak in resistant lines (r = 0.348; Fisher’s z-test, *p* < 0.05). (D) Temporal dynamics of *RpYa7-1* and *RpCCL* abundance in resistant versus susceptible maize tissues following aphid removal. Lines represent mean abundance values; error bars show SEM. Data are separated by plant sites. Resistant, orange; susceptible, blue. (E) Estimated marginal means (emmeans) of log10-transformed *RpYa7-1* and *RpCCL* abundance in resistant and susceptible maize genotypes at FS (0 h post aphid removal). Bar heights indicate emmeans; error bars represent SEM from linear mixed-effects models. Significance in (D) indicates *p* < 0.05; (E) shows *p* values from linear mixed-effects models with Tukey post hoc test.

We next asked whether maize resistance influences the accumulation of aphid-derived RNAs in host tissues. Plants were infested for 24 h, aphids were removed, and tissues were collected from three sites (FS, UFS, and DFS) at 0, 12, and 24 h post-removal. We quantified two aphid RNAs: the *RpYa7-1* (*Ya*) and the mRNA *RpCCL*. In susceptible lines, *RpYa7-1* and *RpCCL* levels were strongly correlated (*r* = 0.734, *p* = 0.015, *n* = 70), consistent with coordinated accumulation (Fig. 6C). In resistant lines, however, this correlation was significantly weaker (*r* = 0.348, *p* = 0.002, *n* = 50), indicating more independent dynamics. A formal comparison of correlation coefficients confirmed that the association was stronger in susceptible lines (*p* < 0.05, Fisher’s *z*-test), suggesting a decoupling of RNA stability in resistant backgrounds.

To further dissect these dynamics, we applied linear mixed-effects models. *RpYa7-1* abundance was unaffected by maize resistance, with no significant effects of genotype, site, or time, and no interactions (all *p* > 0.05, Table S3). Accordingly, *RpYa7-1* remained stable across sites and time points in both resistant and susceptible lines (Fig. 6A). By contrast, *RpCCL* exhibited marked genotype-dependent dynamics. ANOVA revealed significant genotype × time (F_2,229_ = 11.64, *p* < 0.001), genotype × site (F_2,229_ = 10.40, *p* < 0.001), time × site (F_4,229_ = 5.43, *p* < 0.001), and three-way interactions (F_4,229_ = 8.05, *p* < 0.001; Table S3, Fig. S12). Post hoc estimated marginal means (emmeans;Table S4) confirmed that *RpCCL* was already reduced in resistant lines at the FS (0 h; emmean difference = −1.07, *p* = 0.018), whereas *RpYa7-1* showed no such difference (emmean difference = −0.16, *p* = 0.62).

Together, these results reveal decoupled regulation of *Ya* and mRNA: mRNA levels are dynamically modulated by host resistance in a site- and time-specific manner, whereas *Ya* remains stable, consistent with a functional aphid RNA that persists within the host.

## Discussions

Our study provided a comprehensive view of aphid-delivered cross-kingdom RNA dynamics in maize–aphid interactions, highlighting both evolutionary and functional aspects of aphid lncRNAs. We identified 21 and 22 *Ya* genes in *R. padi* and *R. maidis*, respectively, revealing conserved and species-specific evolutionary features compared with *M. persicae*. These genes showed conserved chromosomal localization—residing exclusively on chromosome 2 in *Rhopalosiphum* species versus chromosomes 4 and 6 in *M. persicae*—consistent with lineage-specific genomic rearrangements in aphids (36). Phylogenetic analysis further indicated divergent evolutionary trajectories, with some clusters being species-specific and others conserved across species. Collectively, these findings suggest that *Ya* lncRNA-mediated virulence encompasses both conserved and evolved, species-specific mechanisms.

Multiple *Ya* lncRNAs (9 *RpYa* and 12 *RmYa*) were translocated into maize and migrated systemically to distal feeding sites, independent of their expression levels in aphids. Notably, not all highly expressed *Ya* transcripts entered host tissues, indicating that cross-kingdom activity is selective and likely depends on intrinsic RNA properties rather than transcript abundance, in contrast to mobile RNAs reported in plants (41). This pattern also differs from protein effectors, whose delivery is often correlated with salivary gland expression (42). These observations suggest that RNA-specific determinants— such as secondary structure, RNA–protein complexes, or specialized secretion signals— govern cross-kingdom mobility (43).

Most *Ya* lncRNAs were rapidly degraded in maize over time, with only a subset exhibiting sufficient stability to maintain their function. Using *RpYa7-1* and *RmYa5-6* as examples, we showed that these stable *Ya* act as virulence factors that promote aphid colonization on maize. Interestingly, even the less stable *Ya* lncRNAs were capable of systemic movement within maize and may also contribute to virulence. AUG-mutated versions of *RpYa7-1* and *RmYa5-6* retained the ability to enhance aphid fecundity, consistent with observations for *Ya1* in *M. persicae* (33). This confirms that *Ya* lncRNAs act through their RNA molecules rather than via small peptides. Although the putative ORFs of *Ya* are conserved among aphid species (33), this conservation likely reflects an ancient evolutionary constraint that preserves their non-coding function.

The mechanisms by which *Ya* lncRNAs act in plants remain largely unknown. Plant lncRNAs are known to function through diverse modes, including binding to DNA (44), RNA (45), or protein (46, 47), or by generating small RNAs that silence genes (48). Notably, all of these mechanisms ultimately depend on protein partners to mediate function. Supporting this idea, we recently found that *BSCL1* (BPH Salivary gland Cross-kingdom LncRNAs, 49), another lncRNA secreted by brown planthopper (BPH, *Nilaparvata lugens*) into rice, interacts with histone complexes and displaces them from the promoters of defense-related transcription factor genes, resulting in their transcriptional repression (49). Whether *Ya* lncRNAs act through a similar protein-mediated mechanism remains an open question that warrants further investigation.

Beyond *Ya* lncRNAs, we also detected aphid mRNAs translocating into maize tissues. Such cross-kingdom mRNA exchange is not unique to aphid–plant interactions; for example, dodder haustoria can acquire host mRNAs while simultaneously delivering their own transcripts (50, 51). Yet, whether exogenous mRNAs are translated into proteins within recipient plants remains largely unresolved. In one case, *Arabidopsis thaliana* mRNAs transferred into *Botrytis cinerea* were shown to be translated and to compromise fungal infection (52). By contrast, plants appear to maintain highly efficient RNA surveillance systems that restrict the persistence of foreign mRNAs (37). During viral infection, plants recognize double-stranded RNA replication intermediates and recruit Dicer enzymes for degradation (37). In addition, aberrant single-stranded mRNAs are rapidly eliminated through surveillance pathways that rely on ribosome stalling during translation. Unlike mRNAs, lncRNAs generally contain very short or no open reading frames, making them less prone to ribosome engagement (53–55). This property may allow lncRNAs to evade ribosome-mediated decay, offering a potential explanation for why *Ya* lncRNAs are more stable than mRNAs in plant tissues.

Both aphid-delivered lncRNAs and mRNAs displayed distinct degradation patterns across resistant maize genotypes, indicating that long RNAs are not merely passive carriers of genetic information but active participants in the molecular arms race between aphids and host plants. In resistant maize, aphid mRNAs were selectively destabilized while *Ya* lncRNAs remained stable, revealing a decoupling of RNA dynamics that may underlie quantitative resistance. This differential regulation suggests the existence of an exogenous RNA recognition system in maize, potentially governed by resistance QTLs, and points to aphid RNA stability profiles as predictive markers for screening resistant germplasm.

Although genome-wide association studies (GWAS) are powerful tools for identifying resistance genes (24, 56–58), conventional approaches based on aphid fecundity, in some cases together with defense compound concentrations, are limited by the time-consuming and labor-intensive nature of phenotype scoring (24, 56–58). To overcome these constraints, we propose a “cross-kingdom RNA GWAS” (ckRNA-GWAS) framework, which leverages variation in the accumulation of aphid mRNAs and lncRNAs within host tissues to directly link molecular interactions to maize genotypes. This strategy offers a more efficient and mechanistically informative route to discover resistance loci and enables high-throughput identification of germplasm with enhanced aphid resistance.

Our findings uncover a new layer of maize–aphid interactions, where selective translocation and differential RNA stability determine effector activity, while host-driven mRNA degradation underlies resistance. This work reframes cross-kingdom communication by highlighting long RNAs as active players and provides a mechanistically informed framework (ckRNA-GWAS) for accelerating the discovery of resistance loci in crops.

## Acknowledgments

This project is funded by the National Natural Science Foundation of China (project No. 32172392 to Y.C.), the National Key Research and Development Program of China (project No. 2023YFF1000703 to Y.C.), the Hubei Hongshan Laboratory (project No. 2022hszd026 to Y.C.), the Startup Foundation for Advanced Talents at HZAU to Y.C., the Fundamental Research Funds for the Central Universities (Program No. 2022ZKPY003 to Y.C.), and the Wuhan Yingcai Talent Program to Y.C. We thank Professor Jianbing Yan and Professor Wenqiang Li at Huazhong Agricultural University for providing maize varieties.

## Supplementary Information for

### Materials and Methods

#### Plant growth

Maize seeds were sown in a mixture of peat-based substrate and vermiculite (3:1, v/v) and cultivated in a controlled-environment growth chamber under the following conditions: 22 °C and a 16h/8h light/dark cycle. B73, Mo17, and 20 CUBIC lines (1) were kindly provided by Professor Jianbing Yan at Huazhong Agricultural University.

#### Aphid rearing

The aphid species *R. padi* and *R. maidis* were obtained from the lab of Professor Julian Chen at the Chinese Academy of Agricultural Sciences and maintained on B73 maize seedlings under the same growth chamber conditions. All the aphid stocks were kept on the plants grown at 22°C and a 16/8 h light/dark photoperiod cycle in the growth chamber. Aphid stocks used for all experiments have been maintained in the laboratory for more than two years.

#### Identification of *Ya* genes

Publicly available RNA-seq datasets (PRJNA559238 and PRJNA97450) were used for lncRNA annotation. Reference genomes of *Rhopalosiphum padi* and *Rhopalosiphum maidis* are available in NCBI under accessions GCA_020882245.1 and GCA_003676215.3, respectively.

LncRNA gene identification was performed in three main steps. First, strand-specific RNA-seq reads were mapped to the reference genomes. Raw reads were subjected to quality control and adapter trimming using Fastp v0.23.4 (2) with default parameters (quality threshold: Phred score ≥ 15; minimum read length: 15 bp; automatic adapter detection enabled) to ensure high-quality data for downstream analysis. The processed reads were aligned to the genomes using HISAT2 v2.2.1 (3) with default settings (minimum intron length: 20 bp; maximum intron length: 500,000 bp; RNA strandness:unstranded). The resulting alignments were assembled into transcripts using StringTie v2.2.1 (4) in default mode (minimum transcript abundance: 0.1 FPKM; minimum junction coverage: 1; RNA strandness:unstranded). Transcript sequences were extracted with Gffread v0.12.7 (5) using the parameters -V -H -U -N -P -J -M -K -Q -Y -Z -F --keep-exon-attrs to retain detailed exon structure information.

Secondly, LGC (Long Genomic Region Classifier) v1.0 (6) was used for identifying transcripts with non-coding features based on the relationship between ORF (open reading frame) Length and GC content. Meanwhile, assembled transcripts were subjected to the CPC2 (Coding Potential Calculator 2) (7) to calculate the coding potential. The intersection of the results from LGC and CPC2 was the putative lncRNAs. Finally, the candidate lncRNA transcripts were screened against the Rfam v14.3(8) database using default parameters (E-value ≤ 1e-5) to remove known housekeeping RNAs, including rRNAs, tRNAs, snoRNAs, and other structured RNA families.

To annotate the *Ya* gene family in the genomes of *R. padi* and *R. maidis,* the sequence of *M. persicae Ya*1 (*MpYa1*) was used as a query, as described in Chen et al (2020) (9). BLASTn was performed to align *MpYa1* against the annotated lncRNA transcripts with an E-value cutoff of 1e-5 and a similarity threshold of 80%. This analysis identified 21 *Ya* genes in *R. padi* (*RpYa*) and 22 in *R. maidis* (*RmYa*).

#### Genome synteny analyses

For the genome synteny analysis, the 1:1:1 orthologs among *R. padi*, *R. maidis*, and *M. perisicae* genomes were extracted from OrthoFinder’s (10) result and fed to MCScanX_h (11) with BLASTP (E-value ≤ 1e⁻⁵), which was used with “-b 2” option to get the inter-species collinearity among *R. padi*, *R. maidis*, and *M. perisicae*. SynVisio (12) was used to visualize the genome synteny.

#### Aphid feeding experiments to analyze the migration of *Ya* lncRNAs in maize plants and the stability of aphid-delivered RNAs

50 aphids were confined on the fourth leaf of 4 leaf stage B73 maize plants (15 days) using clip cages for a 48-hour feeding period. Samples collected included the aphids themselves, the feeding site (FS), the up-feeding site (UFS), and the down-feeding site (DFS). Counterparts on maize plants without aphid treatment were harvested as controls. All tissue samples from both groups were washed three times with sterilized water, rapidly frozen in liquid nitrogen, and stored at −80°C. 4 replicated per sites were harvested. Those samples were used for RNA extraction for RT-PCR and RNA-seq.

Similar to the feeding experiments described above, 50 aphids were caged on the FS for 48 hours, then removed. Tissue samples were collected from FS, UFS, and DFS at 0 h, 12 h, and 24 h post-insect removal. 4 replicates were harvested for each time point and tissue type. Those samples were subjected to RNA extraction for further analysis of the aphid-delivered RNA stability.

Experiments on seven maize genotypes (B73, Mo17, E28, Ji853, Zi330, H21, and Nx110) were conducted in same way. Each genotype-time point-tissue combination included 4 replicates. Total RNA was extracted from each sample, reverse-transcribed, and subjected to RT-PCR analysis using gene-specific primers (Table S1) to measure transcript abundance.

#### Translocation efficiency accumulation and analysis the stability

To quantify the translocation efficiency of *Ya* RNAs, transcript abundances were measured in both aphids and FS tissues for each candidate gene. *Ya* RNA levels in aphids were normalized to housekeeping genes (*RpRpL7* for *R. padi* and *RmEF1α* for *R. maidis*), while *Ya* RNA levels in maize tissues were normalized to *ZmActin1*. For each replicate and gene, translocation efficiency was defined as the ratio of transcript abundance in FS to that in aphids (FS/Aphids). To avoid division by zero, a pseudo-value of 10^6 was assigned when transcripts were absent in aphids but present in FS. The resulting efficiency matrix was summarized by calculating the mean, standard deviation (SD), and coefficient of variation (CV) for each gene (13). Statistical differences in translocation efficiency among genes were evaluated using the Kruskal–Wallis test, followed by Dunn’s post hoc test with Bonferroni correction. Significance levels were indicated as *\*p* < 0.05, \*\**p* < 0.01, and \*\*\**p* < 0.001.

To assess the stability of aphid-delivered RNAs, transcript abundances of selected *Ya* RNAs were monitored in maize tissues at 0, 12, and 24 h after aphid removal. RNA levels were normalized to *ZmActin1*, and stability was estimated by calculating the CV of transcript abundance across biological replicates for both *R. maidis* and *R. padi*.

#### Sequencing of 3’ ends of *RpYa7-1* and *RmYa5-6* transcripts

To obtain the full-length cDNA of *RpYa7-1* and *RmYa5-6*, 3’ rapid amplification of cDNA ends (RACE) was performed. For 3’ RACE, 3 μg of aphid total RNA was added to an 80 μL ligation mixture containing the following: 4 μL T4 RNA ligase, 8 μL 10× T4 RNA ligase buffer, and 8 μL BSA (all from the T4 RNA Ligase kit, Thermo Fisher Scientific, EL0021, USA); 8 μL ATP (Thermo Fisher Scientific, R0441, USA); and 10 pmol of 3’ RACE RNA adaptor (Table S1). Ligation was carried out overnight at 16 °C. The ligated RNA was reverse-transcribed into cDNA using an oligonucleotide complementary to the 3’ RACE adaptor with the RevertAid First Strand cDNA Synthesis Kit (Thermo Fisher Scientific, K1622, USA).

RACE PCRs were performed using gene-specific primers (*RpYa7-1-*RACE-F and *RmYa5-6-*RACE-F as forward primers) in combination with primers corresponding to the oligo sequence complementary to the 3′ RNA adapter (Table S1). Each 50 μL PCR reaction contained 1 μL Phanta® Max Super-Fidelity DNA Polymerase (Vazyme, P505, Nanjing, China), 25 μL Phanta Max Buffer, 1 μL dNTP mix, 2 μL cDNA template, 2 μL of each primer (10 μM), and 17 μL nuclease-free water. The cycling program was: 95 °C for 3 min, followed by 35 cycles of 95 °C for 15 s, 55-62 °C (corresponding annealing temperature for each lane is shown in Fig. S7A)for 15 s, and 72 °C for 20 s, with a final extension at 72 °C for 5 min. PCR products were purified, ligated into the pEASY®-Blunt Cloning vector (TransGen,CB101,Beijing,China), and sequenced to confirm the full-length cDNA of *RpYa7-1* and *RmYa5-6*.

#### Subcellular Localization of *RpYa7-1* and *RmYa5-6* lncRNAs in *N. benthamiana* cells

For visualization of *RpYa7-1* and *RmYa5-6* in plant cells, an RNA-triggered fluorescence (RTF) reporter system was employed (14). The RTF system consists of two modules: (i) an RNA switch containing a probe sequence complementary to the target RNA, and (ii) a GFP expression cassette that is regulated by the RNA switch. In the absence of the target RNA, the RNA switch remains in an inactive conformation, leading to recruitment of the 26S proteasome and degradation of GFP, thereby preventing fluorescence. In contrast, binding of the target RNA to the probe region induces a conformational switch that prevents GFP degradation, allowing GFP accumulation and fluorescence detection.

The probe sequence specific for *RpYa7-1* and *RmYa5-6* was designed to avoid secondary structure interference and off-target hybridization and was cloned into the pC1300-AtU6::AptRS-AtUbq10::mSunTag vector at the designated restriction sites, generating the RNA-switch RTF construct (Fig. S7). Primers used for cloning are listed in Table S1. The construct was sequence-verified by Sanger sequencing and introduced into *A. tumefaciens* GV3101(pSoup-p19) (Weidi, AC1003, Beijing, China) by heat shock. The RTF construct was co-infiltrated with pC1300-AtUbq10::RpYa7-1 or pC1300-AtUbq10::RmYa5-6 into fully expanded leaves of 4-week-old *N. benthamiana* plants using the leaf infiltration method. *A. tumefaciens* cultures were grown overnight in LB medium containing appropriate antibiotics, harvested by centrifugation, and resuspended in infiltration buffer (100 mM MgCl₂, 10 mM MES pH 5.6, 400 µM acetosyringone) to an OD₆₀₀ of 0.6. Equal volumes of GV3101(pSoup-p19) strains carrying the respective constructs were mixed prior to infiltration. Infiltration was performed on the abaxial side of leaves using a 1-mL needleless syringe. Plants infiltrated with only the RTF construct or pC1300-AtUbi10::RpYa7-1 or pC1300-AtUbi10::RmYa5-6 served as negative controls, while those infiltrated with two constructs for GFP expression.

2 days post-infiltration (dpi), infiltrated leaves were harvested, mounted in distilled water on glass slides, and imaged using a Leica SP8 confocal laser scanning microscope (Leica Microsystems, Germany). GFP fluorescence was excited at 488 nm and detected between 500–530 nm. Chlorophyll autofluorescence was collected at 650–670 nm to assist with subcellular localization. Laser power and detector settings were kept constant across all samples to allow comparative analysis. Images were processed using ImageJ software (National Institutes of Health, USA).

#### RNA interference

Double-stranded RNAs (dsRNAs) targeting the *RpYa7-1* and *RmYa5-6* genes were synthesized and purified in vitro using the T7 High Yield RNA Transcription Kit (Vazyme, TR101-01, Nanjing, China) according to the manufacturer’s instructions. The gene-specific primers used for dsRNA synthesis are listed in Table S1. The artificial diet (15) was prepared by dissolving the synthesized dsRNA in a sucrose solution at final concentrations of 0.75% (w/v) for *R. padi* and 1% (w/v) for *R. maidis*, with the addition of 0.02% (w/v) Neutral Red dye (Sigma-Aldrich, C1022, USA) as a feeding tracer. Third-instar nymphs were starved for 2 hours before the experiment. Groups of 20 aphids were then transferred to feeding chambers and allowed to feed on the dsRNA-sucrose solution for 48 hours through a Parafilm membrane. 8 independent biological replicates were set up for each dsRNA treatment group. After the 48-hour feeding period, individuals showing red coloration in the abdomen were collected for subsequent gene expression and phenotypic analysis. Total RNA was extracted, and the transcript levels of the target genes were quantified by RT-PCR to evaluate the efficiency of gene silencing. The specific primers used for RT-PCR are listed in Table S1. Aphids fed on a diet containing ds*LTA* (16) were used as the negative control.

#### Virus-mediated transient expression of lncRNA

To investigate the effects of *RpYa7-1* and *RmYa5-6* on the colonization of two aphid species, we employed the Foxtail mosaic virus (FoMV) plasmid PV101-EGFP(17) to express lncRNAs in maize. FoMV is a positive-sense, single-stranded RNA virus with a genome of approximately 6.2 kb that contains five open reading frames (ORFs) encoding proteins such as replicase and the capsid protein (CP). The enhanced green EGFP gene and lncRNA were inserted between ORF4, which encodes the movement protein, and the CP gene. This arrangement allows both EGFP and lncRNA to be integrated into the viral genome, ensuring their replication and expression during viral infection. EGFP fluorescence provides a visual marker for viral expression and localization. cDNA synthesized from total RNA of *R. padi* and *R. maidis* was used to amplify the transcripts of *RpYa7-1* and *RmYa5-6*, respectively. The recombinant plasmids PV101-EGFP-U6::RpYa7-1 and PV101-EGFP-U6::RmYa5-6 were constructed by inserting *RpYa7-1* and *RmYa5-6*, respectively, together with the U6 promoter(U6 promoter was amplified using the universal forward primer U6 promoter-F, and the reverse primers PV101-EGFP-U6::RpYa7-1 and PV101-EGFP-U6::RmYa5-6 corresponded to U6_*RpYa7-1*-R and U6_*RmYa5-6*-R, respectively) into PV101-EGFP using the ClonExpress Ultra One Step Cloning Kit (Vazyme,C115, Nanjing, China). In addition, a non-coding fragment of the EGFP gene (76–495 nt) was amplified using primers tGFP-F/tGFP-R with PV101-EGFP as the template to serve as a negative control, hereafter referred to as *tGFP*. The U6 promoter and *tGFP* fragment were inserted together into PV101-EGFP to generate the plasmid PV101-EGFP-U6::tGFP, with the U6 promoter amplified using primers U6 promoter-F and U6_tGFP-R.

To assess whether *RpYa7-1* and *RmYa5-6* encode small peptides, we used the NCBI ORF Finder (standard genetic code; minimum ORF length, 10 amino acids) (https://www.ncbi.nlm.nih.gov/orffinder/). This analysis identified a predominant open reading frame in each transcript: one beginning at nucleotide 211 (AUG) in *RpYa7-1* and another at nucleotide 266 (AUG) in *RmYa5-6*.

To determine whether the putative small peptide encoded by the *Ya* gene influences aphid colonization, mutant plasmids were generated by site-directed mutagenesis of the predicted start codons. For the *RpYa7-1* mutant (PV101-EGFP-U6::RpYa7-1-mut), the AUG codon at position 211 was replaced with a TAG stop codon. Using PV101-EGFP-U6::Rp7-1 as a template, two fragments were amplified: Fragment 1 with primers *RpYa7-1*-F1/*RpYa7-1*-R1 and Fragment 2 with primers *RpYa7-1*-F2/*RpYa7-1*-R2. These fragments were subsequently used as templates for a secondary PCR with bridging primers (*RpYa7-1*-F/*RpYa7-1*-R1-L and *RpYa7-1*-F2-L/*RpYa7-1*-R), producing 250 bp and 276 bp products, respectively. The two fragments and U6 promoter (amplified using primers U6_*RpYa7-1*-R and U6 promoter-F) were assembled into the linearized PV101-EGFP-U6::RpYa7-1 backbone using the ClonExpress Ultra One Step Cloning Kit (Vazyme, C115, Nanjing, China). A similar strategy was applied to construct PV101-EGFP-U6::RmYa5-6-mut, in which the ATG codon at position 266 in *RmYa5-6* was replaced with TAG. Fragments 1 and 2 were amplified from PV101-EGFP-U6::RmYa5-6 using primer pairs *Rm5-6*-F1/*Rm5-6*-R1 and *Rm5-6*-F2/*Rm5-6*-R2, then re-amplified with bridging primers (*RmYa5-6*-F/*RmYa5-6*-R1-L and *RmYa5-6*-F2-L/*RmYa5-6*-R) to yield 195 bp and 275 bp products. These fragments, together with the U6 promoter (amplified using primers U6_*RmYa5-6*-R and U6 promoter-F), were assembled into PV101-EGFP-U6::RmYa5-6 using the same kit. Sequences of all primers used for constructing FoMV plasmids are listed in Table S1. All plasmids were introduced into *A. tumefaciens* strain GV3101 (pSoup-p19) (Weidi, AC1003, Beijing, China) via heat shock and infiltrated into fully expanded leaves of 4-week-old *N.benthamiana* plants using the leaf infiltration method. *A. tumefaciens* cultures were grown overnight in LB medium supplemented with the appropriate antibiotics, harvested by centrifugation, and resuspended in infiltration buffer (100 mM MgCl₂, 10 mM MES, pH 5.6, and 400 µM acetosyringone) to an OD₆₀₀ of 0.6. Infiltration was performed on the abaxial side of leaves using a 1mL needleless syringe. For inoculation of B73 maize plants, sap was extracted from *N. benthamiana* leaves agroinfiltrated as described above and harvested at 3 dpi. Leaves were ground in 0.67% (w/v) deionized water using a mortar and pestle, and the resulting sap was supplemented with 1% (w/v) Celite 545 AW (Sigma-Aldrich, 22141, USA) for rub inoculation of the first three leaves of 3-leaf-stage maize seedlings. Five minutes or more after inoculation, leaves were rinsed with tap water to remove residual sap and Celite. Plants were then bagged and maintained under high humidity and low light for approximately 24 h before being returned to standard growth conditions. Each construct was typically inoculated into at least 12 maize plants per experiment, which were subsequently subjected to aphid bioassays at 14 dpi, when strong GFP fluorescence was observed.

#### Aphid fecundity assays and honeydew secretion assay

To assess the impact of gene silencing on aphid reproduction, a single aphid that had been fed on the target dsRNA (or ds*LTA* control) diet for 48 hours was transferred onto a B73 maize plant at the 4 leaf stage. The number of nymphs produced by each aphid was recorded daily for 10 consecutive days. This experiment included 10 replicates per treatment group.

The effect of gene silencing on aphid feeding behavior was evaluated by measuring honeydew excretion. 5 aphids from the respective dsRNA-treated groups were transferred onto a 4 leaf stage B73 maize seedling. The seedling was placed in a customized setup, with a piece of filter paper positioned beneath the plant to collect honeydew droplets. After 48 hours, the filter paper was carefully removed and stained with a water-soluble aniline blue solution (18) to visualize the honeydew spots. The total stained area on each filter paper, which correlates with the volume of honeydew excreted, was quantified using ImageJ software (National Institutes of Health, USA). 10 replicates were performed for each group, with the ds*LTA*-treated group serving as the control.

To evaluate the impact of overexpression plants on aphid reproduction, 1 to 2 adult aphids were inoculated onto each overexpression maize plant. After 24 hours, all adult aphids and excess nymphs were carefully removed, leaving exactly 2 nymphs per plant. These nymphs were allowed to develop into adults. Upon reaching adulthood, the number of nymphs produced per adult per day was recorded daily for a consecutive period of 10 days. The experiment included 10 replicates per plant group.

To assess the effect of overexpression plants on aphid feeding behavior, a similar inoculation procedure was followed. One to two adult aphids were placed on each plant and removed after 24 hours, again leaving two nymphs per plant. After these nymphs matured into adults, their feeding activity was quantified by measuring honeydew excretion. Adults were placed in the setup depicted in Fig. S9A, where honeydew was collected on a filter paper for 48 hours. The filter paper was subsequently stained with a water-soluble aniline blue solution to visualize the honeydew spots. The total stained area, which correlates with the amount of honeydew produced, was quantified using ImageJ software (National Institutes of Health, USA). This experiment was also performed with 10 replicates per group.

#### RNA extraction, cDNA synthesis and RT-PCR

All plant and aphid samples collected during the experiments were rapidly frozen in liquid nitrogen and stored at −80°C. Total RNA was extracted from collected tissues using TRIzol reagent (Invitrogen, 15596018, USA) and treated with RNase-free DNase I (Thermo Fisher Scientific, EN0521, USA) to remove residual genomic DNA. First-strand cDNA was synthesized from 1 μg of total RNA using a mixture of oligo(dT) and random primers with the HiScript II 1st Strand cDNA Synthesis Kit (Vazyme, R211, Nanjing, China), following the manufacturer’s instructions. Quantitative RT-PCR was performed on a CFX96 Real-Time System (Bio-Rad, USA) using ChamQ Blue Universal SYBR qPCR Master Mix (Vazyme, Q312, Nanjing, China). The primer sequences for aphid *Ya* lncRNAs and mRNAs are listed in Table S1. For evaluating aphid silencing efficiency and quantifying the expression levels of *Ya* lncRNAs in insects, *RpRPL7* or *RmEF1α* was used as the reference gene. For all other experiments involving plant materials, *ZmActin1* was selected as the internal reference gene. Relative expression was calculated using the 2^−ΔCt^ method and log10-transformed (with a 10^-10^ constant to avoid log(0) issues) for analysis.

#### RNA library preparation and sequencing

RNA-seq was performed by Novogene Co., Ltd. (Beijing, China). Samples were frozen in liquid nitrogen, ground to a fine powder using sterilized stainless-steel beads in a TissueLyser II (Qiagen, Germany), and homogenized in TRIzol reagent (Invitrogen, 15596018, Waltham, MA, USA) following the manufacturer’s protocol. RNA concentration and purity were measured with a NanoDrop 2000 spectrophotometer (Thermo Scientific, USA), and RNA integrity was assessed with an Agilent 2100 Bioanalyzer (Agilent Technologies, USA). Quantification was performed using a Qubit 2.0 Fluorometer (Thermo Scientific, USA). Only samples with RNA integrity number (RIN) >7.0 were used for library construction.

For each sample, 1 μg of high-quality total RNA was used to prepare strand-specific libraries with the VAHTS^®^ Universal V6 RNA-seq Library Prep Kit (Vazyme, NR604, Nanjing,China). The libraries were assessed for quality and fragment size distribution using the Bioanalyzer 2100. Sequencing was performed on an Illumina NovaSeq 6000 platform (Illumina,USA), generating 150-bp paired-end reads. RNA-seq data deposited at NCBI under project number PRJNA1313817.

#### Identification of aphid-delivered RNAs in FS on maize by RNA-seq analysis

FS samples and the controls were used for analysis of the aphid-delivered RNAs in FS on maize. The genome of B73 (Assembly ID: GCA_902167145.1) merged with *R. padi* genome was used for analysis of maize FS samples by *R. padi* (Assembly ID: GCA_020882245.1), genome of B73 merged with *R. maidis* genome for analysis of maize FS samples by *R. maidis* (Assembly ID: GCA_003676215.3).

Raw sequencing reads were processed with fastp (v0.20.1) (2) and adapters and low-quality reads (phred quality score <25) were removed. Reads shorter than 50 bp after trimming were discarded. The quality of clean reads was evaluated using FastQC (v0.11.9) (19). Only clean reads passing quality control were retained for downstream analysis. Clean reads were aligned to the reference genome using RMTA (v2.6.3) with the HISAT2 aligner (3). Parameters were set to trim 15 bases from the 5′ end of each read, and minimum and maximum intron lengths were specified as 20 and 500,000, respectively. The resulting BAM files were used for read quantification with HTSeq (v0.6.1) (20), using the parameters -r, -i gene_id, -t exon, and library-specific strand settings (-s no for non-stranded or -s yes for stranded data). Only uniquely mapped reads overlapping annotated exons were counted. Transcript abundance was also quantified as TPM (transcripts per million) using TPMCalculator (v0.0.3) (21). BAM files generated by RMTA and the corresponding GTF annotation files were provided to TPMCalculator (-b for BAM, -g for annotation). Strand-specificity was set according to the RNA library type (--stranded no for non-stranded libraries).

#### Aphid fecundity on different maize varieties

20 maize varieties were randomly chosen from the CUBIC population. To evaluate the impact of these varieties on the fecundity of two aphid species, a standardized bioassay was conducted. For each genotype, plants at the 4 leaf stage were used. One to two apterous adult aphids were carefully transferred onto each plant. After 24 hours, all adults and excess nymphs were removed, leaving exactly 1 nymphs per plant to standardize the age and density of the test cohort. These nymphs were allowed to develop into adults under controlled environmental conditions. Upon reaching adulthood, the number of nymphs produced per adult per day was recorded daily for 10 consecutive days. The experiment included four biological replicates per maize genotype–aphid species combination.

Five CUBIC varieties (E28, Ji853, Zi330, H21, and Nx110), which exhibited significant impact on aphid fecundity, along with B73 and Mo17, were repeated for the assay, with 10 replicates per maize genotype.

#### Classification and mixed-effects modeling analysis

Analyses were conducted in R (version 4.3.1). Genotypes were classified using linear models (*lm*) of nymph count by genotype, with B73 as the reference (susceptible). Genotypes with significantly lower nymph counts (*p* < 0.05, negative estimate) were classified as resistant; others were susceptible. Expression data were analyzed using linear mixed-effects models (lme, nlme package) for log10-transformed *Ya* and mRNA expression, with fixed effects of Trait (resistant or susceptible), Time (0, 12, 24 hours), Site (FS, UFS, DFS), and their interactions, and a random effect of Genotype. *ANOVA* was used to test main effects and interactions. Estimated marginal means (emmeans, emmeans package) and pairwise contrasts (Resistant vs. Susceptible) were computed for each Time and Site combination, with Time 0, FS contrasts adjusted to reported values (*Ya*: *p* = 0.62; mRNA: *p* = 0.018). Results were visualized using ggplot2 as line plots, a bar plot for Time 0, and emmeans bar plots. *ANOVA* results and emmeans were listed in the Table S4.

#### Data analysis

Data analyses were performed in R and GraphPad Prism 10 software (GraphPad Software, USA). All statistical tests are described in the figure legends.

---

**Figure S1.**
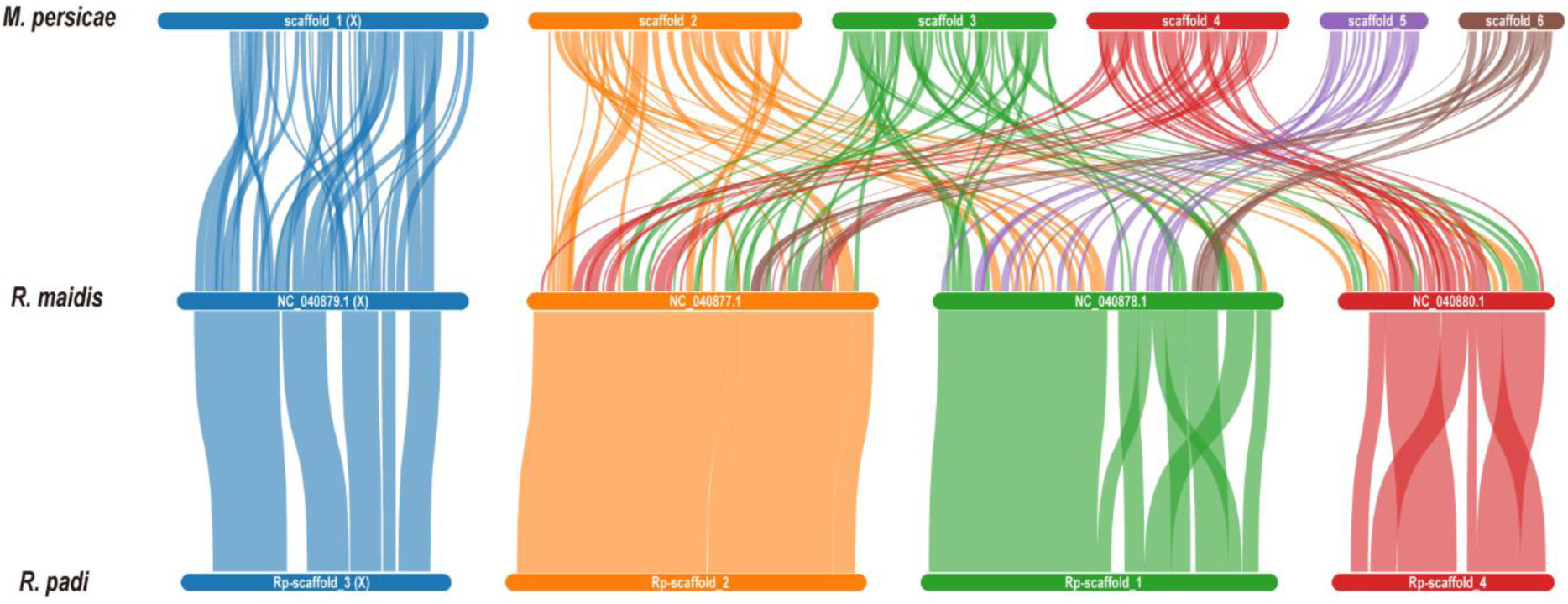
Synteny analysis of *M. persicae*, *R. maidis* and *R. padi.* Colored lines indicate collinear blocks, highlighting conserved genomic regions and structural variations among the three genomes.

**Figure S2.**
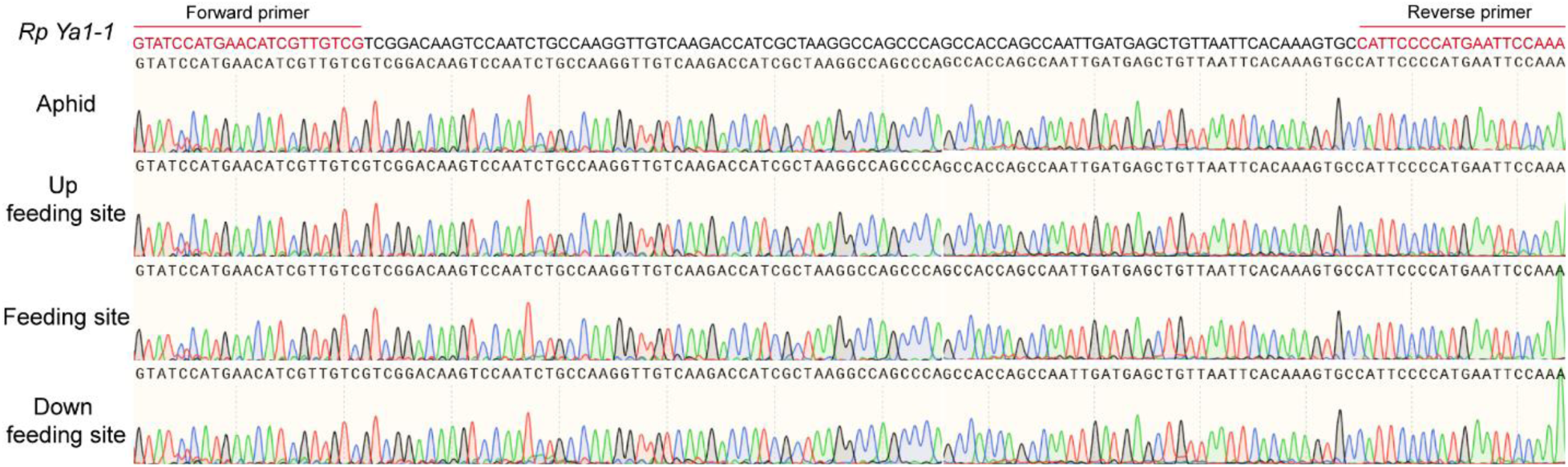
Sequence verification of *RpYa1-1* in aphid and maize tissues. *RpYa1-1* transcripts were amplified from aphid, feeding site (FS), up feeding site (UFS), and up feeding site (DFS) by RT-PCR, and the PCR products were sequenced and aligned to the reference sequence to confirm transcript identity. Primer sequences used for amplification are listed in Table S1. Sequence alignment shows the nucleotide-level consistency between the amplified products and the annotated *RpYa1-1* reference sequence, validating the presence of the transcript in both aphid and maize tissues.

**Figure S3.**
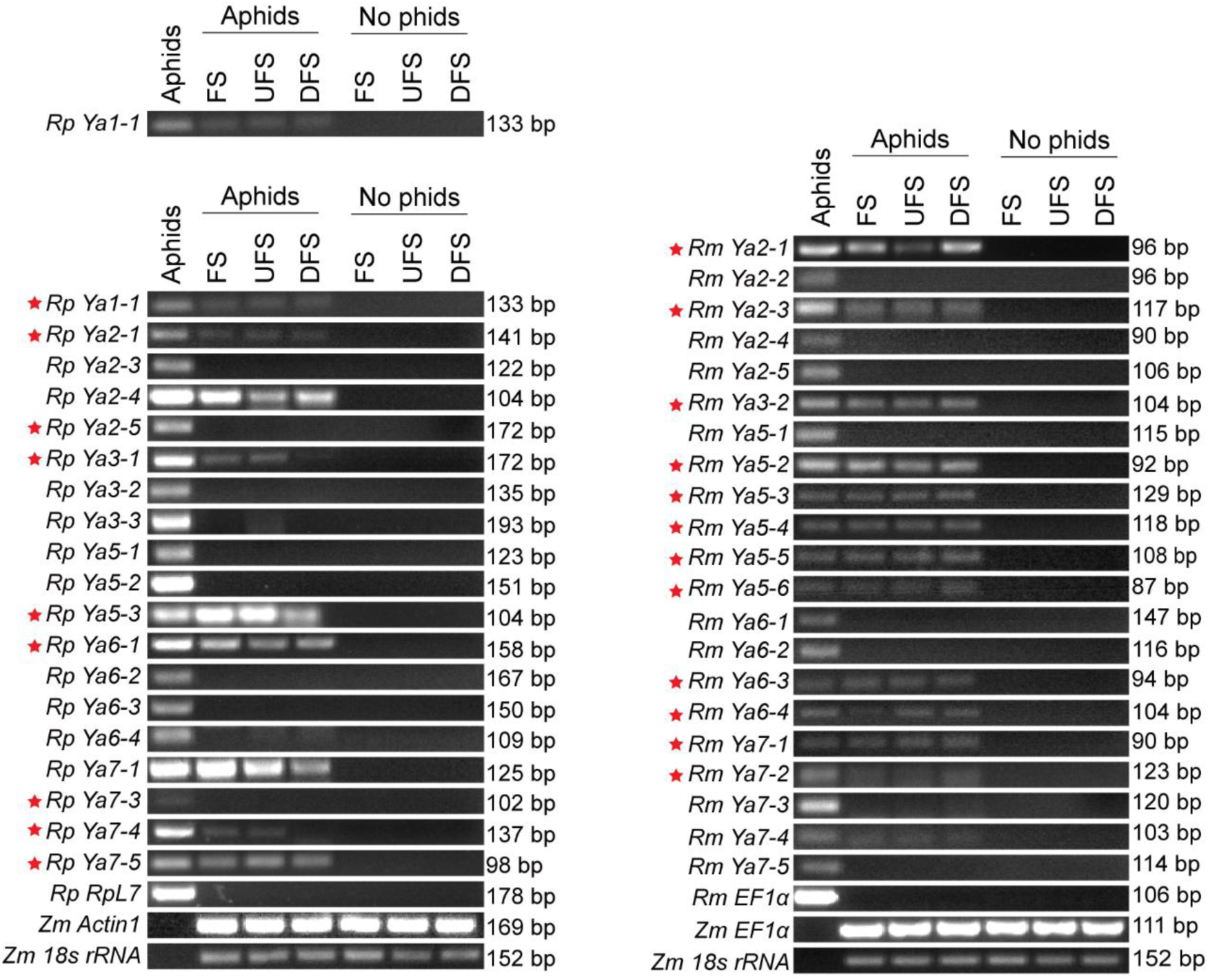
Systemic migration of 9 *RpYa* and 12 *RmYa* transcripts in maize. *Ya* transcripts capable of transfer and systemic migration in maize are indicated with red stars. The right side of the gel shows the sizes of the PCR products. Experimental setup is shown in Fig. 2A. The *Ya* transcripts were amplified by RT-PCR using gene-specific primers (Table S1) and PCR products were separated by 2% agarose gel. The bands corresponding to the PCR products were sequenced directly by forward and/or reverse primers to verify the identity of the *Ya* sequence.

**Figure S4.**
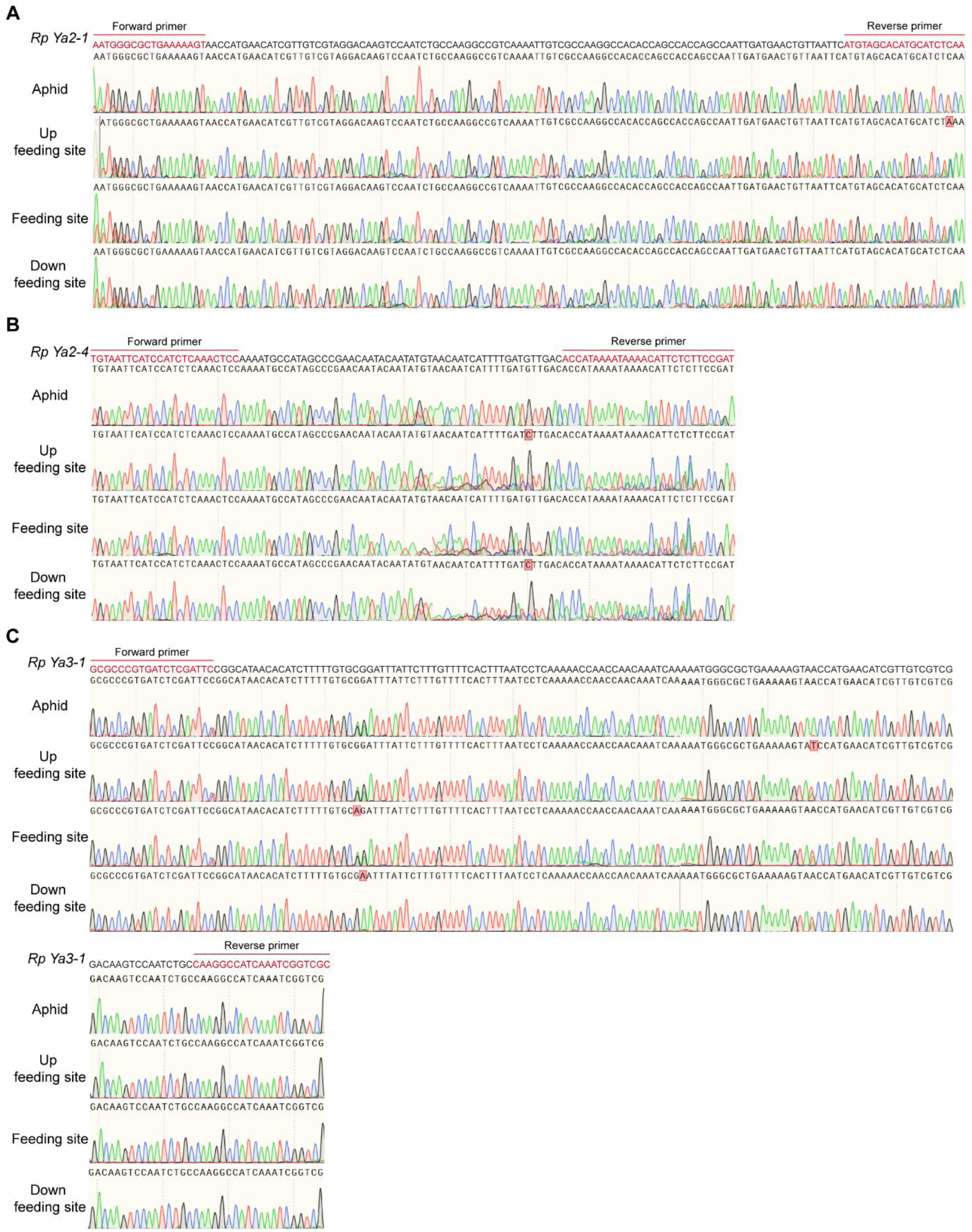
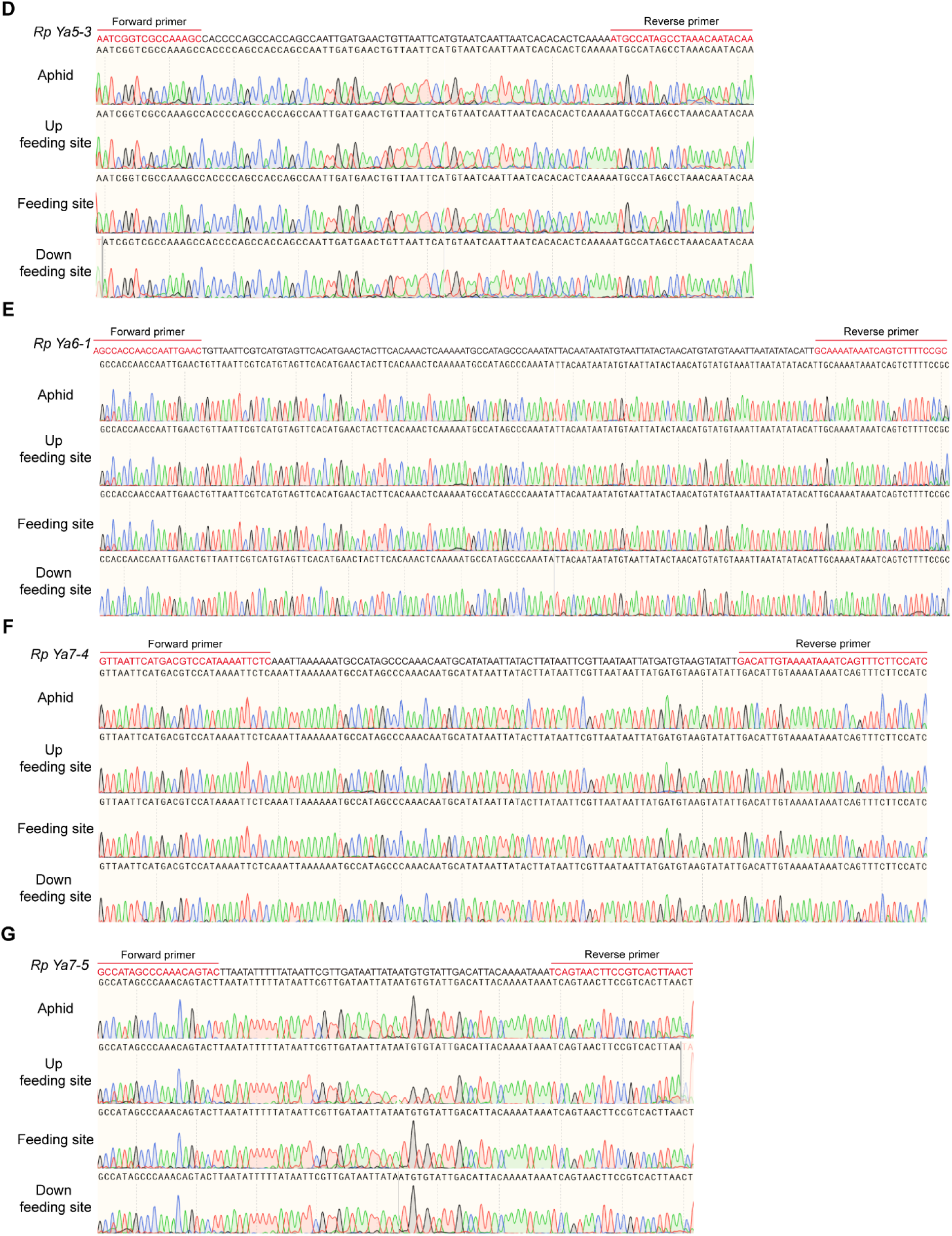
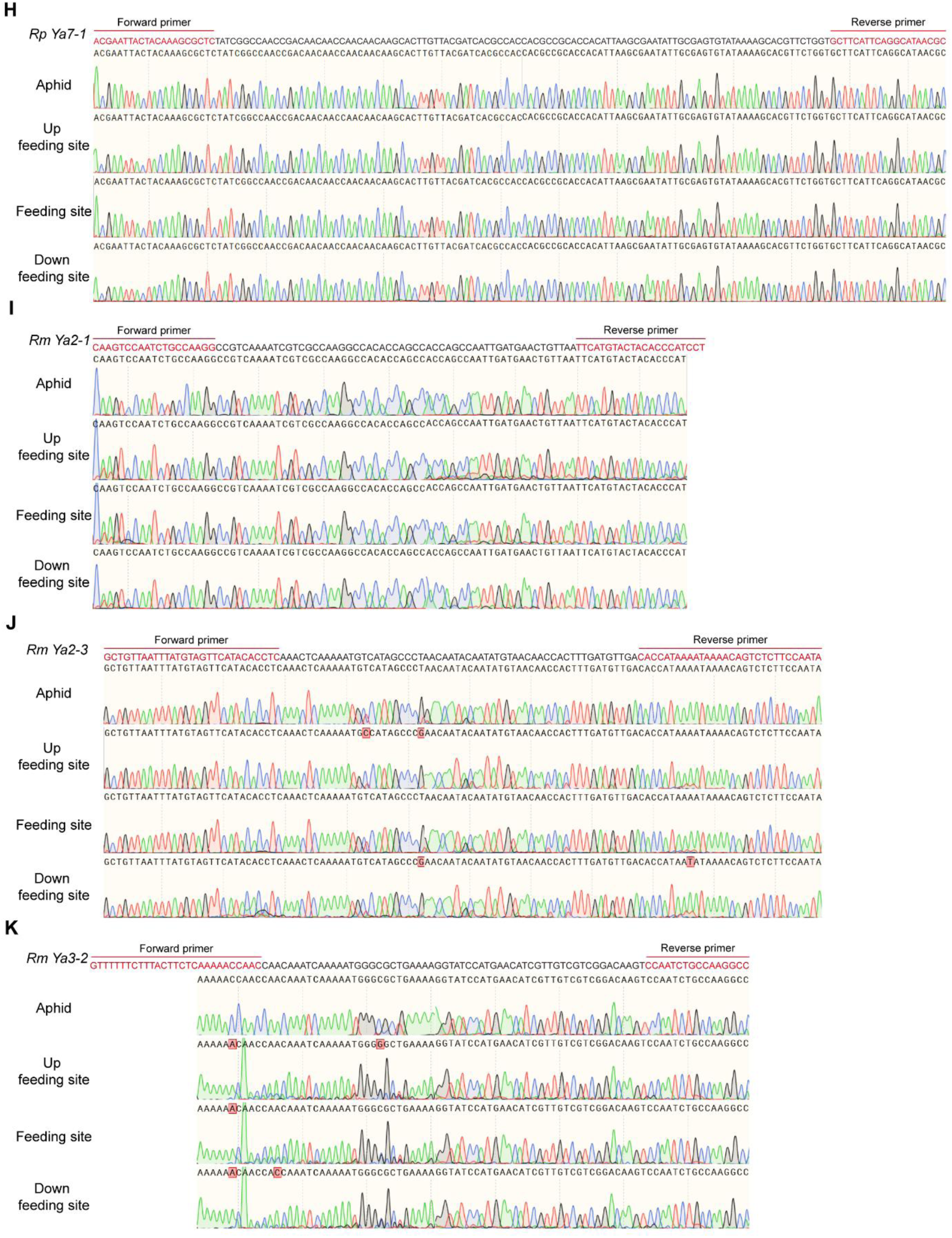
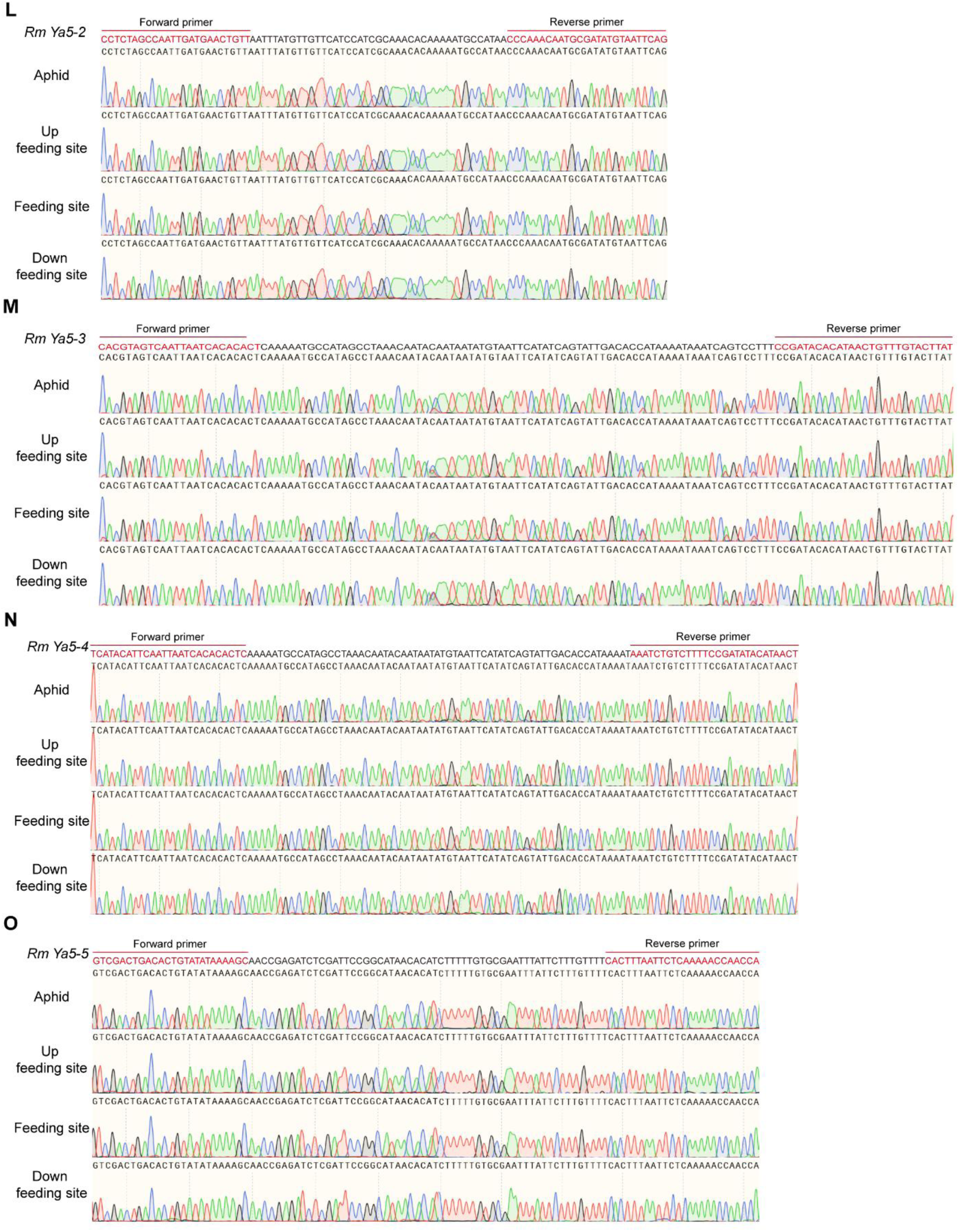
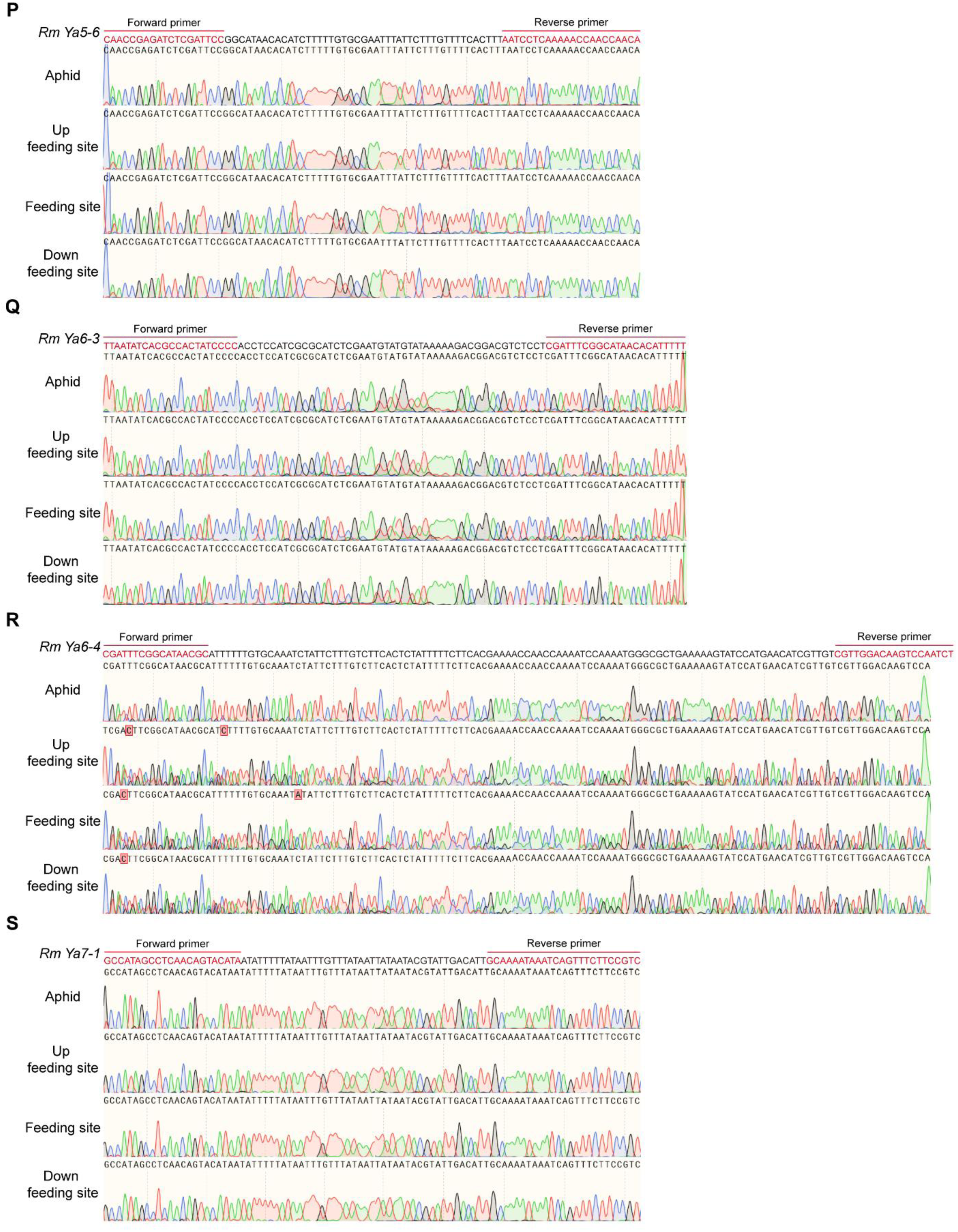
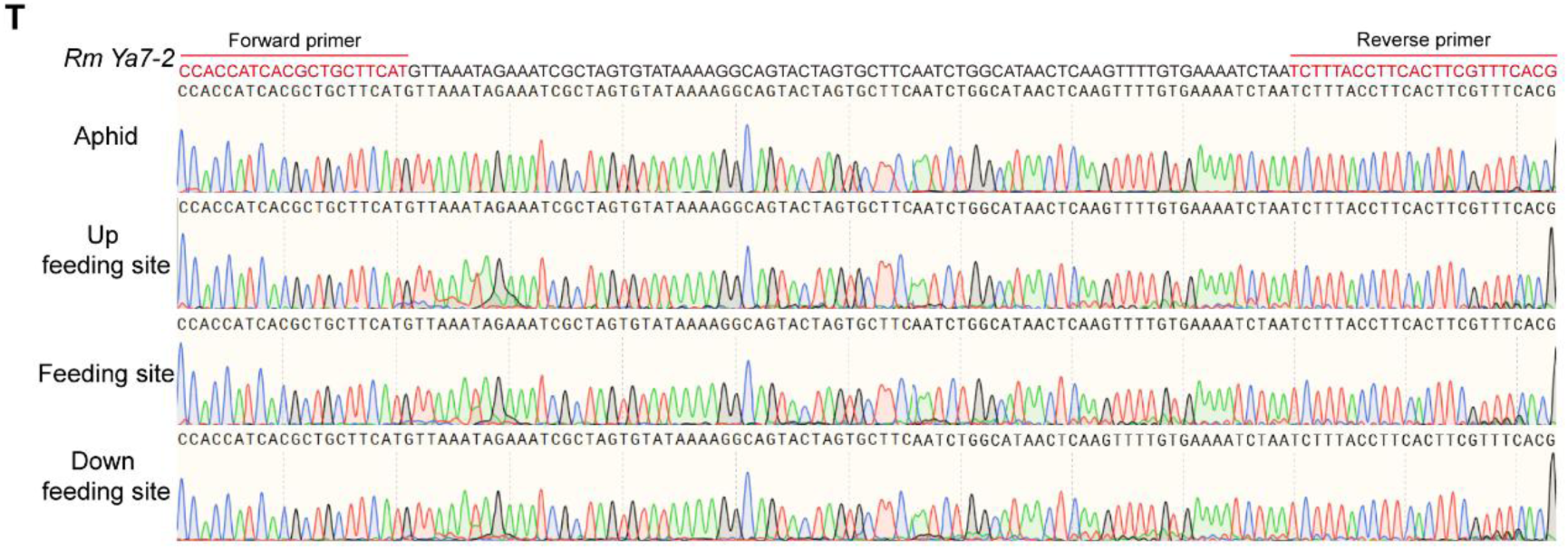
Sequence verification of cross-kingdom *Ya* transcripts in aphid and maize tissues. **A–H** show cross-kingdom *RpYa* lncRNAs (except *RpYa1-1*). **I–T** show cross-kingdom *RmYa* lncRNAs. Cross-kingdom *Ya* transcripts were amplified from aphid, feeding site, up feeding site, and down feeding site by RT-PCR, and the PCR products were sequenced and aligned to the reference sequence to confirm transcript identity. Primer sequences used for amplification are listed in Table S1. Sequence alignment shows the nucleotide-level consistency between the amplified products and the annotated these transcripts reference sequences, validating the presence of these transcripts in both aphid and maize tissues.

**Figure S5.**
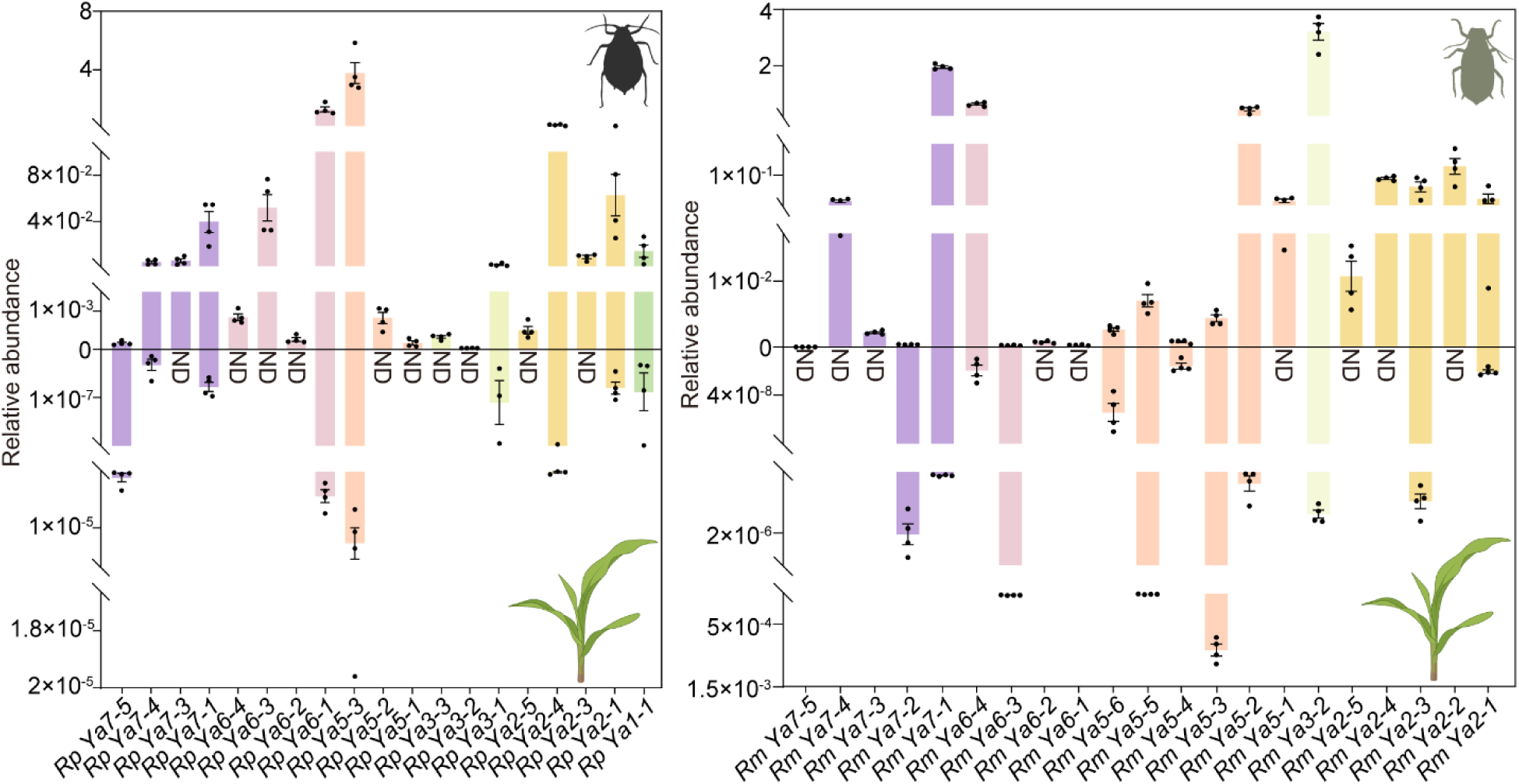
Relative abundance of *Ya* transcripts in aphids and FS. Relative abundance of *RpYa* (left) and *RmYa* (right) transcripts in aphids and maize FS. Upper panels show relative abundance in aphids; lower panels show relative abundance at FS. Experimental setup is shown in Fig. 2A. Relative abundance was measured by RT-PCR with gene-specific primers (Table S1) and normalized to *RpRpL7* (*R. padi*), *RmEF1α* (*R. maidis*), or *ZmActin1* (FS). Points: individual replicates (n = 4); bars: mean ± SD; ND: not detected.

**Figure S6.**
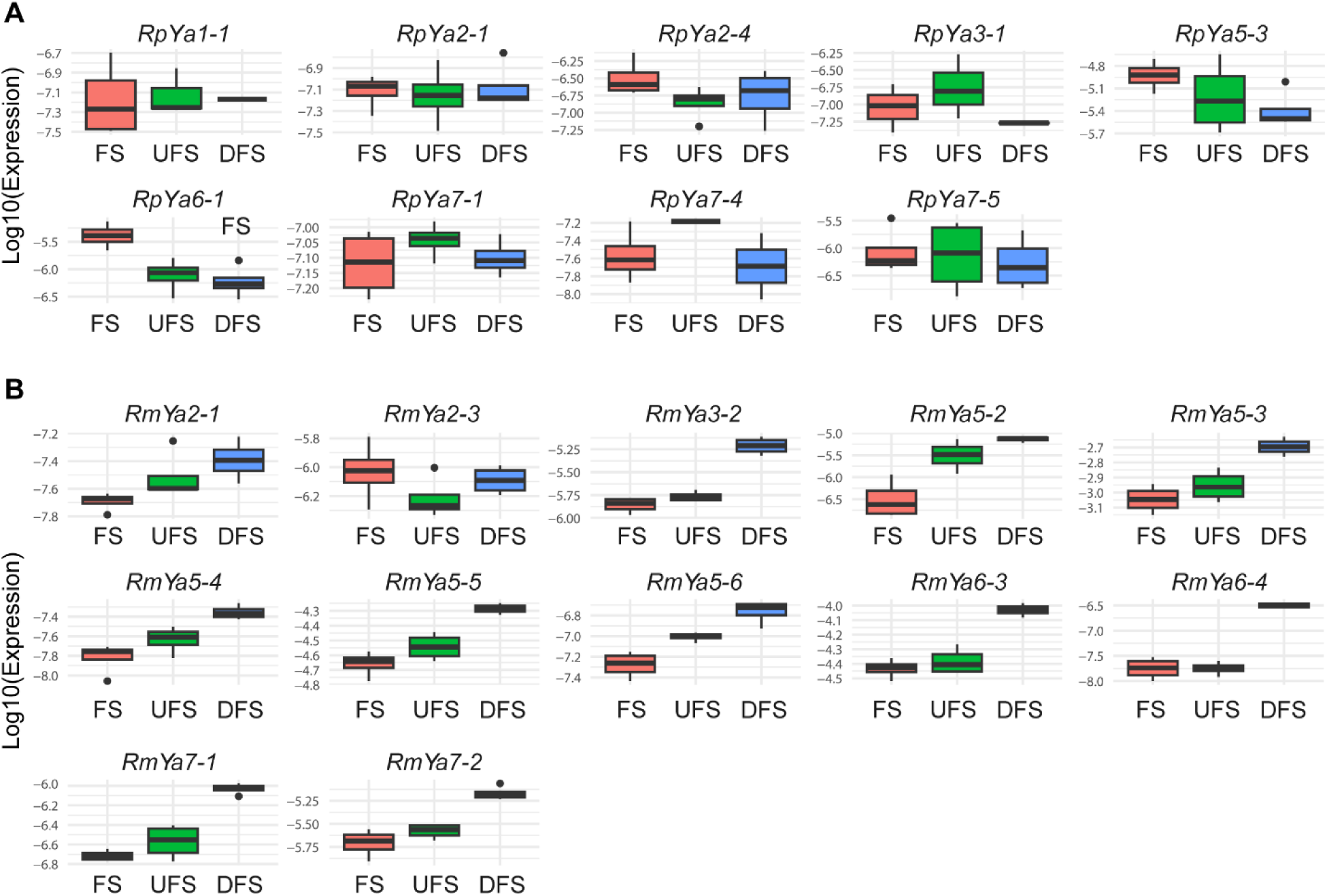
Relative abundance of cross-kingdom *Ya* transcripts in FS、UFS and DFS. Relative abundance of cross-kingdom *RpYa* (A) and *RmYa* (B) transcripts in FS, UFS, and DFS. Experimental setup is shown in Fig. 2A. Boxplots depict log₁₀-transformed *Ya* transcript levels measured by RT-PCR with gene-specific primers (Table S1) and normalized to *ZmActin1*. Boxes represent the interquartile range (Q1–Q3), the central line indicates the median, and whiskers show the minimum and maximum of four biological replicates. The corresponding heatmap is shown in Fig. 2C.

**Figure S7.**
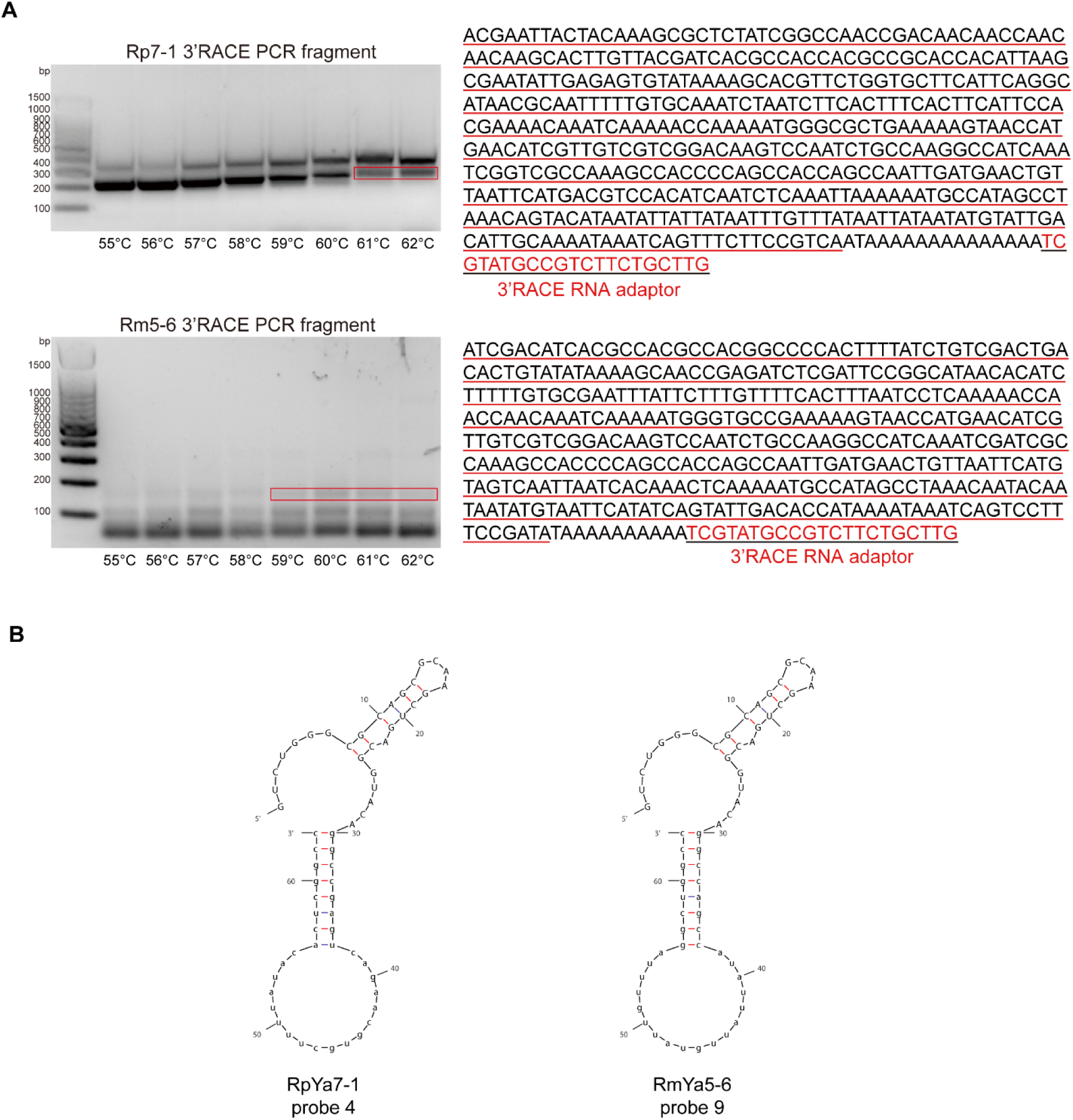
3′RACE experiment and predicted RNA-switch secondary structures of *RpYa7-1* and *RmYa5-6*. (A) Results of a 3’ RACE experiment. The upper panel shows *RpYa7-1* and the lower panel shows *RmYa5-6*. PCR products were separated on a 2% agarose gel; the target bands (highlighted by red boxes) were excised, purified, and sequenced. Lane annotations indicate the corresponding annealing temperatures. The sequences obtained from 3′RACE are shown on the right. Red nucleotides at both ends represent the 3′ RNA adapter sequences, while the underlined red sequence corresponds to the annotated sequence. The downstream sequence was obtained through 3′RACE. (B) Predicted secondary structures of the RNA-switches. The left panel shows the RNA-switch designed for *RpYa7-1*, and the right panel shows the RNA-switch designed for *RmYa5-6*. The design principles and screening criteria of the RNA-switches are described in the *Materials and Methods* section.

**Figure S8.**
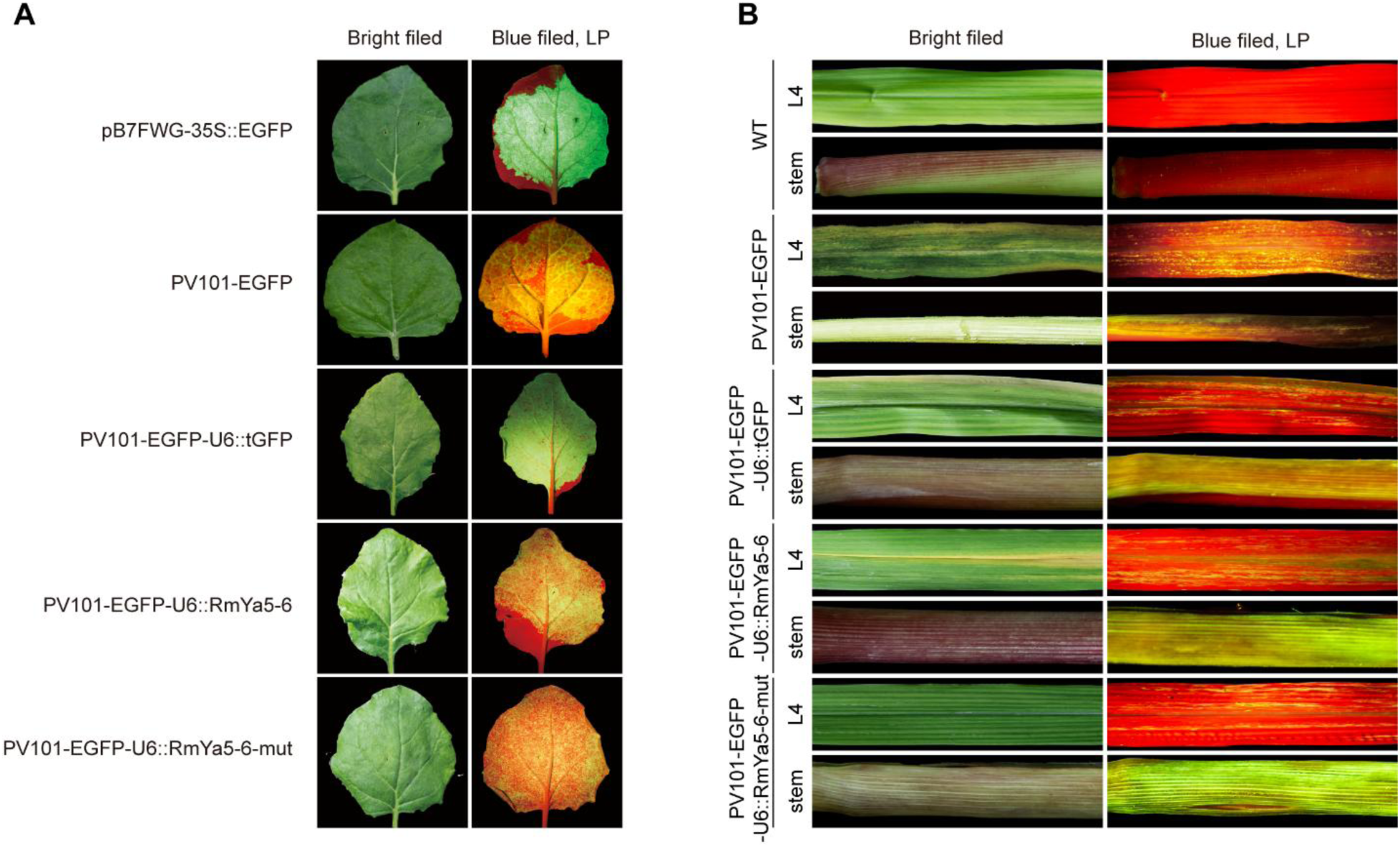
EGFP fluorescence in plants infiltrated with FoMV–lncRNA constructs. (A) EGFP fluorescence in *N.benthamiana* leaves infiltrated with FoMV plasmids expressing different lncRNAs. pB7WG2-35S::EGFP was used as a positive control. (B) EGFP fluorescence in maize stem and the fourth leaf(L4) at 14 dpi. Wild-type (WT) maize was used as a negative control. Fluorescence was observed a 488 nm UV LED light source. Left panels show leaves under normal light, and right panels show EGFP fluorescence under 488 nm excitation.

**Figure S9.**
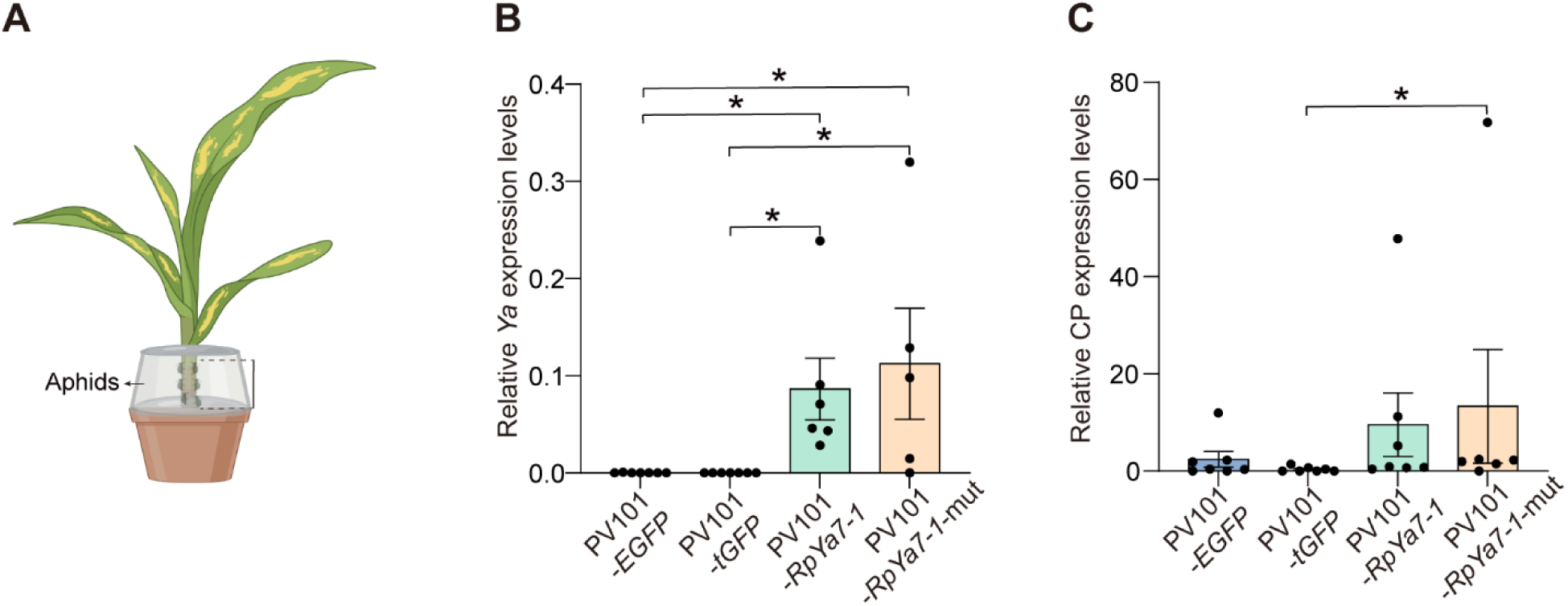
Bioassay setup and expression levels of *Ya* and coat protein expression (CP) in maize inoculated with FoMV–lncRNA constructs. (A) Schematic of the bioassay setup: maize plants at 14 dpi were used for aphid bioassays. Aphids were confined to maize stems with transparent plastic cups, aluminum foil, and cotton to measure fecundity and honeydew production. (B and C) Expression of *Ya* (B) and *CP* genes (C) in maize at 14 days post-inoculation with FoMV–lncRNA constructs. Transcript levels were measured by RT-PCR with gene-specific primers (Table S1) and normalized to *ZmActin1*.. Significance was determined by unpaired two-tailed t-tests: \**p* < 0.05; \*\**p* < 0.01; \*\*\**p* < 0.001; \*\*\*\**p* < 0.0001.

**Figure S10.**
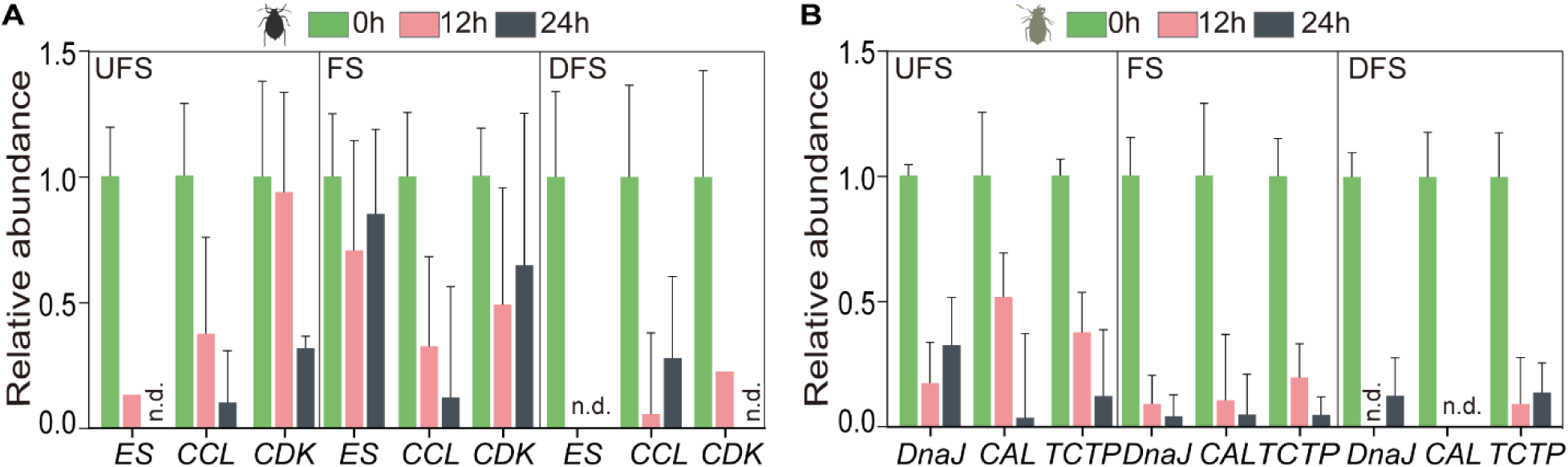
Relative abundance of cross-kingdom aphid mRNAs in FS, UFS and DFS. (A) Relative abundance of cross-kingdom mRNAs from *R. padi* detected in FS, UFS, and DFS: *ES*, *Extended Synaptotagmin-like protein 2*; *CDK*, *Cyclin-dependent Kinase*; *CCL*, *Chemokine (C-C motif) Ligand*. (B) Relative abundance of cross-kingdom mRNAs from *R. maidis* detected in FS, UFS, and DFS: *CAL*, calreticulin; *TCTP*, *Translationally Controlled Tumor Protein*; *DnaJ*, *DnaJ heat shock protein*. Relative abundance was measured by RT-PCR with gene-specific primers (Table S1) and normalized to *ZmActin1*.

**Figure S11.**
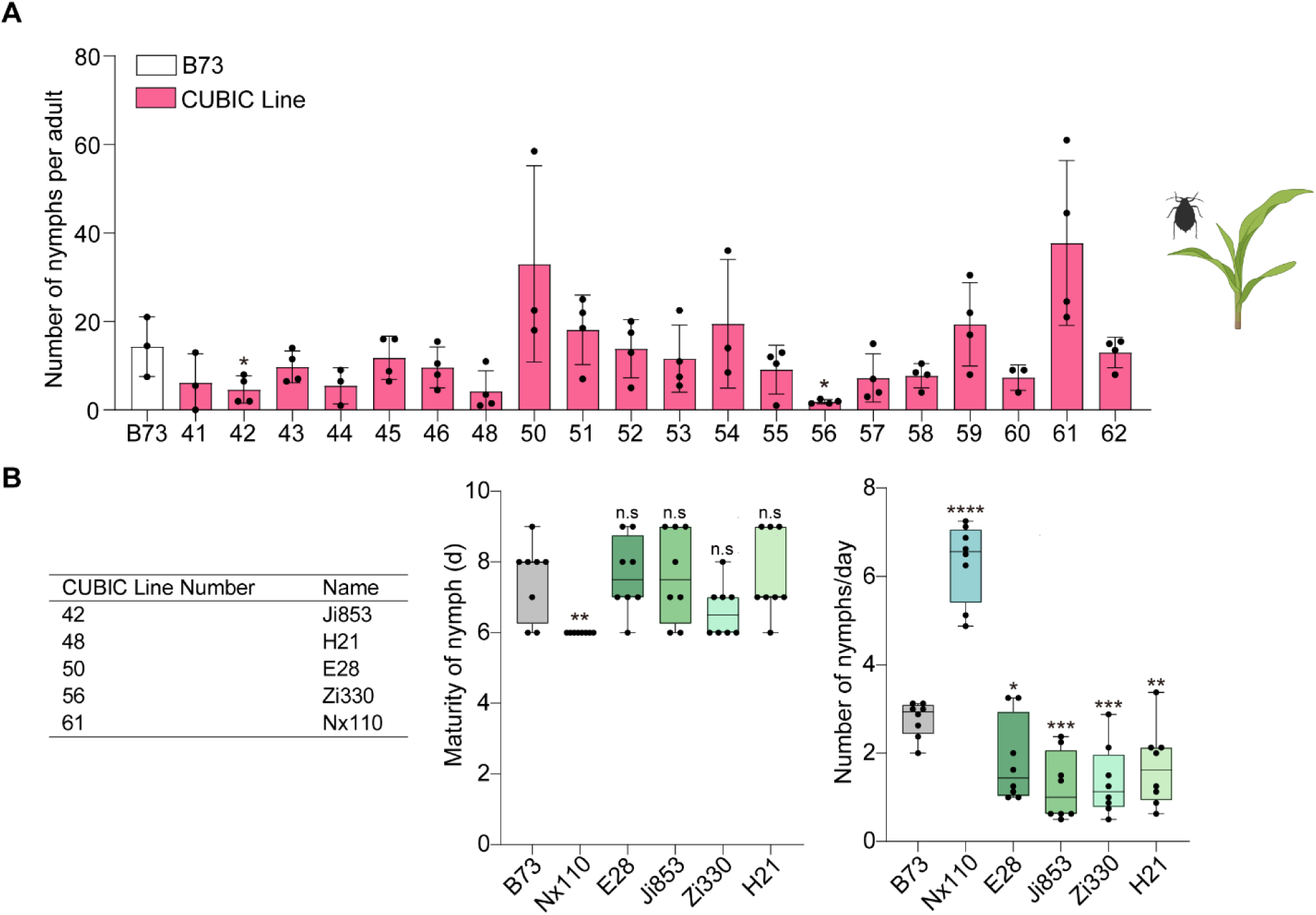
Reproduction and development of *R.padi* on maize CUBIC lines. (A) Reproduction of R. padi on 20 maize CUBIC lines. Each dot represents the daily offspring number from a single aphid (n=3–4 plants per variety). (B) Nymph-to-adult development time and reproduction of *R. padi* on 4 resistant and 1 susceptible maize lines selected from the 20 lines. Each dot represents the daily offspring of a single aphid or its development time to adult (n = 7–8 plants per variety). Significance was determined by unpaired two-tailed t-tests against B73: ns, not significant; \**p* < 0.05; \*\**p* < 0.01; \*\*\**p* < 0.001; \*\*\*\**p* < 0.0001.

**Table S1.**
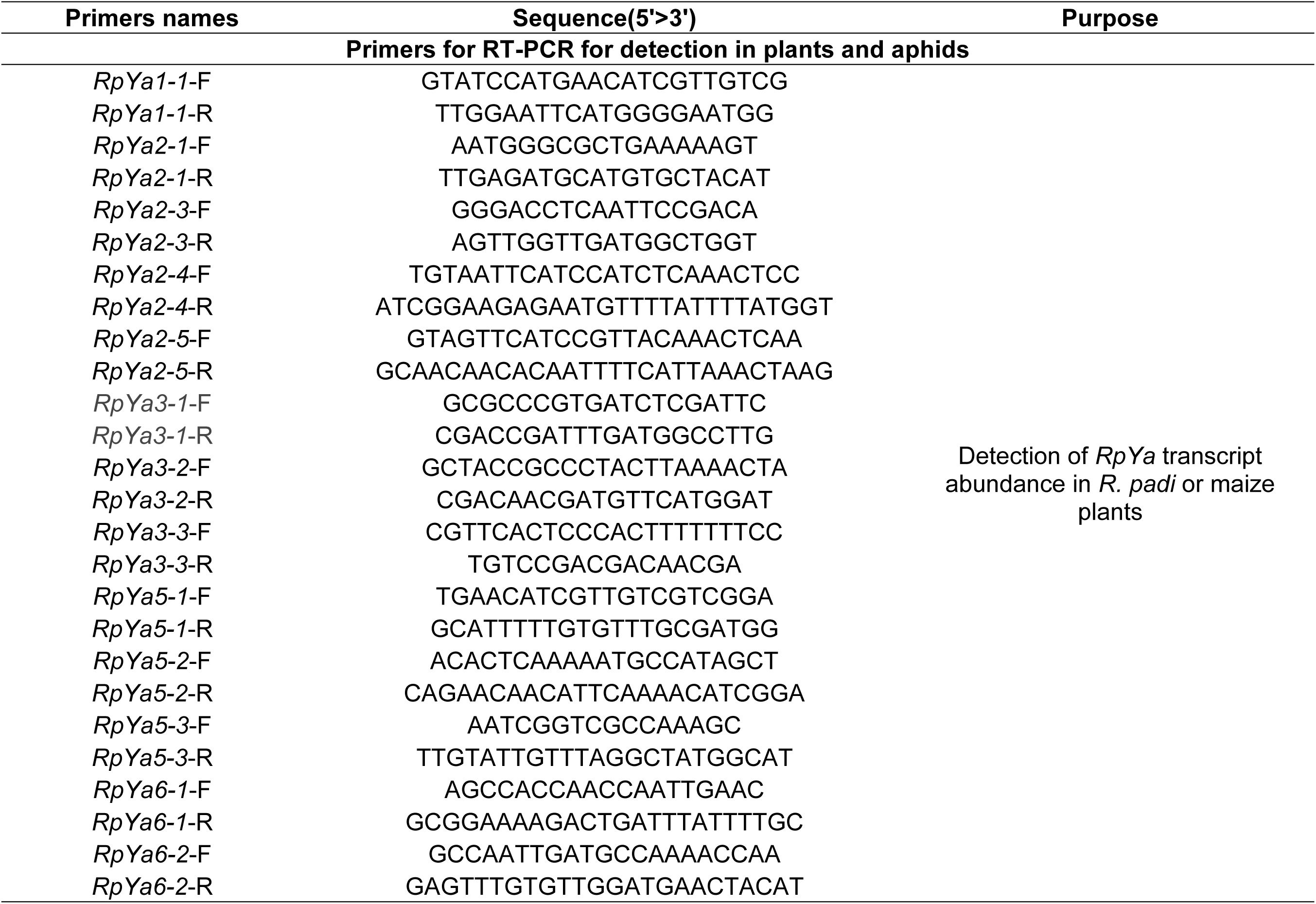
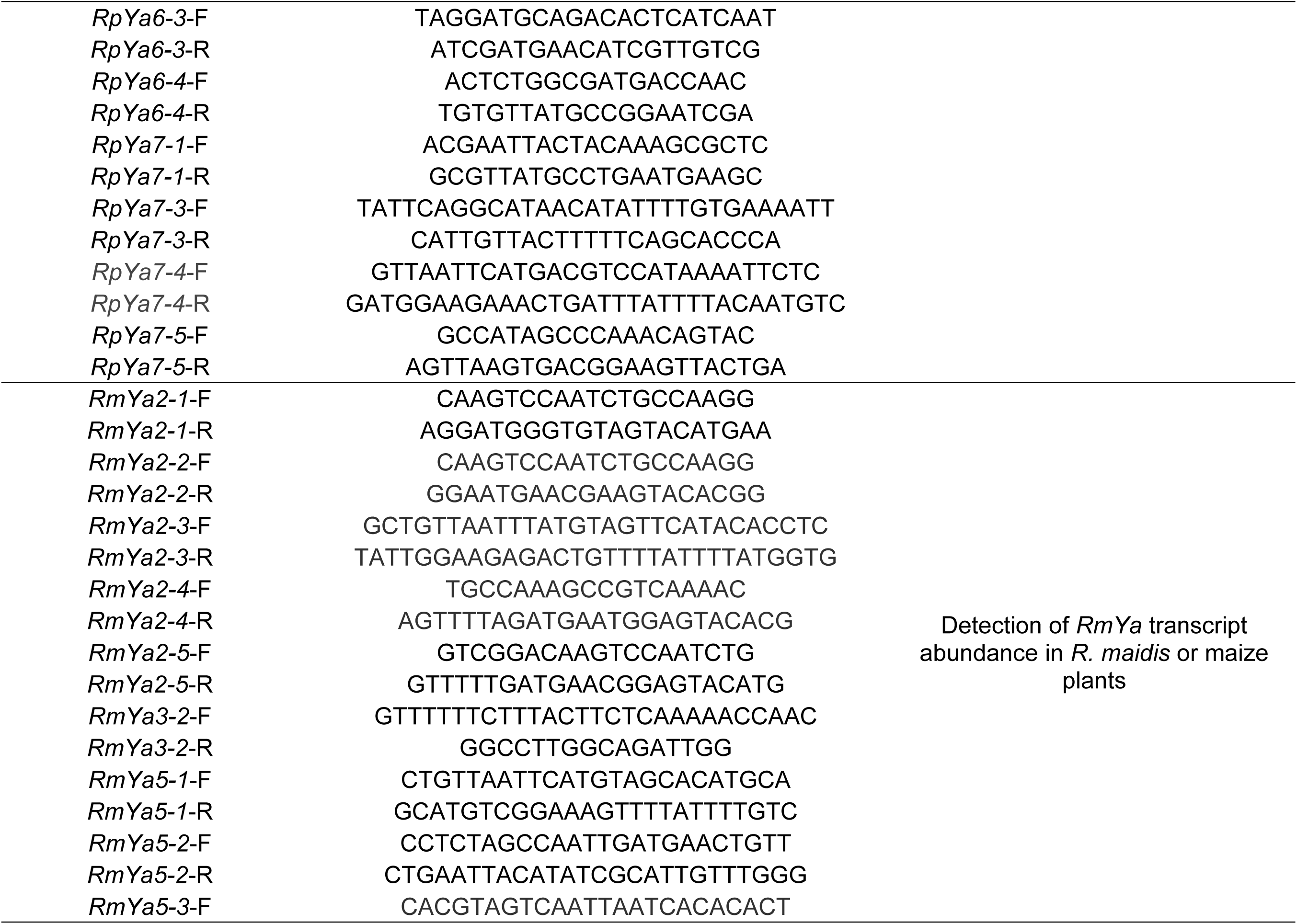
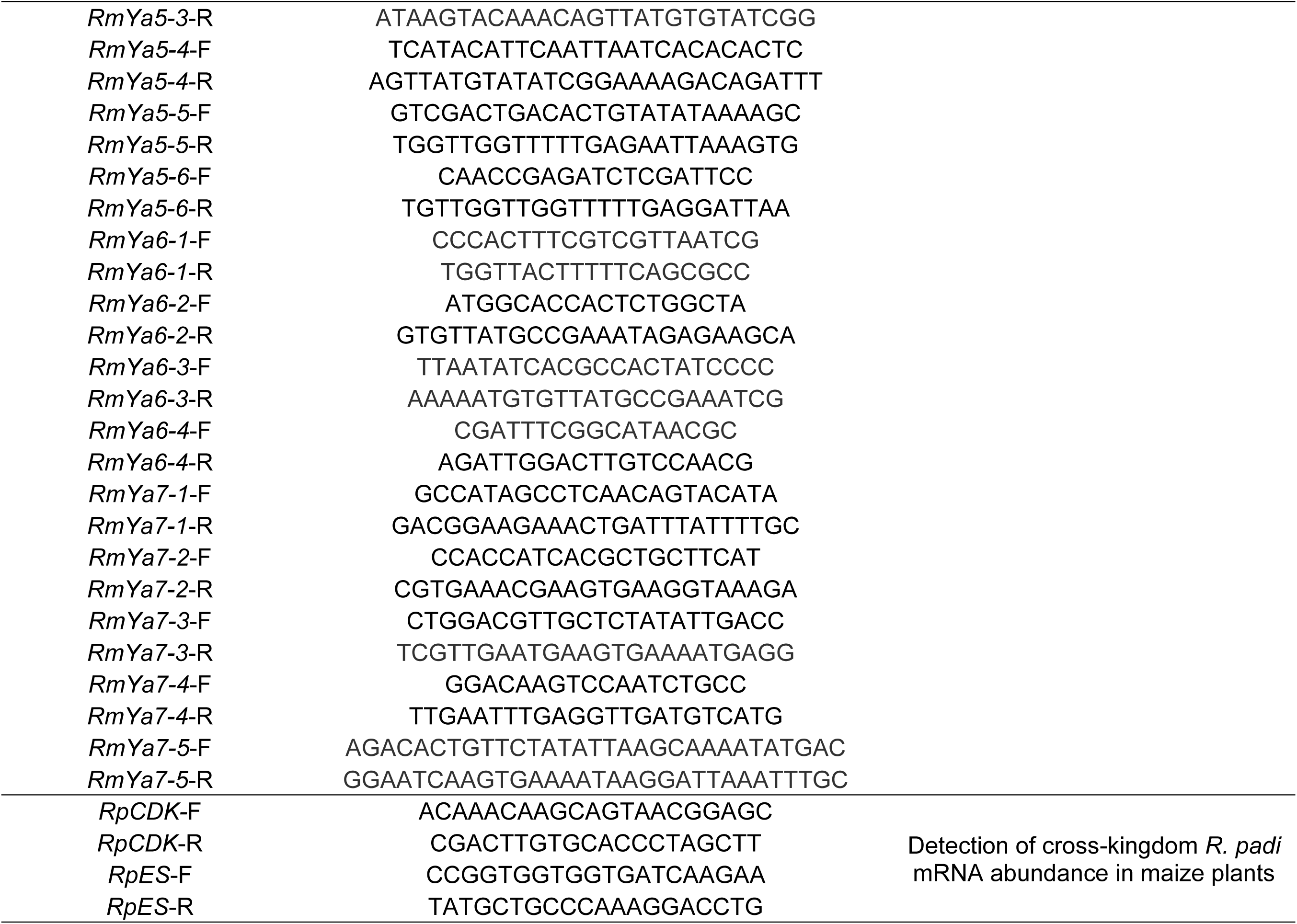
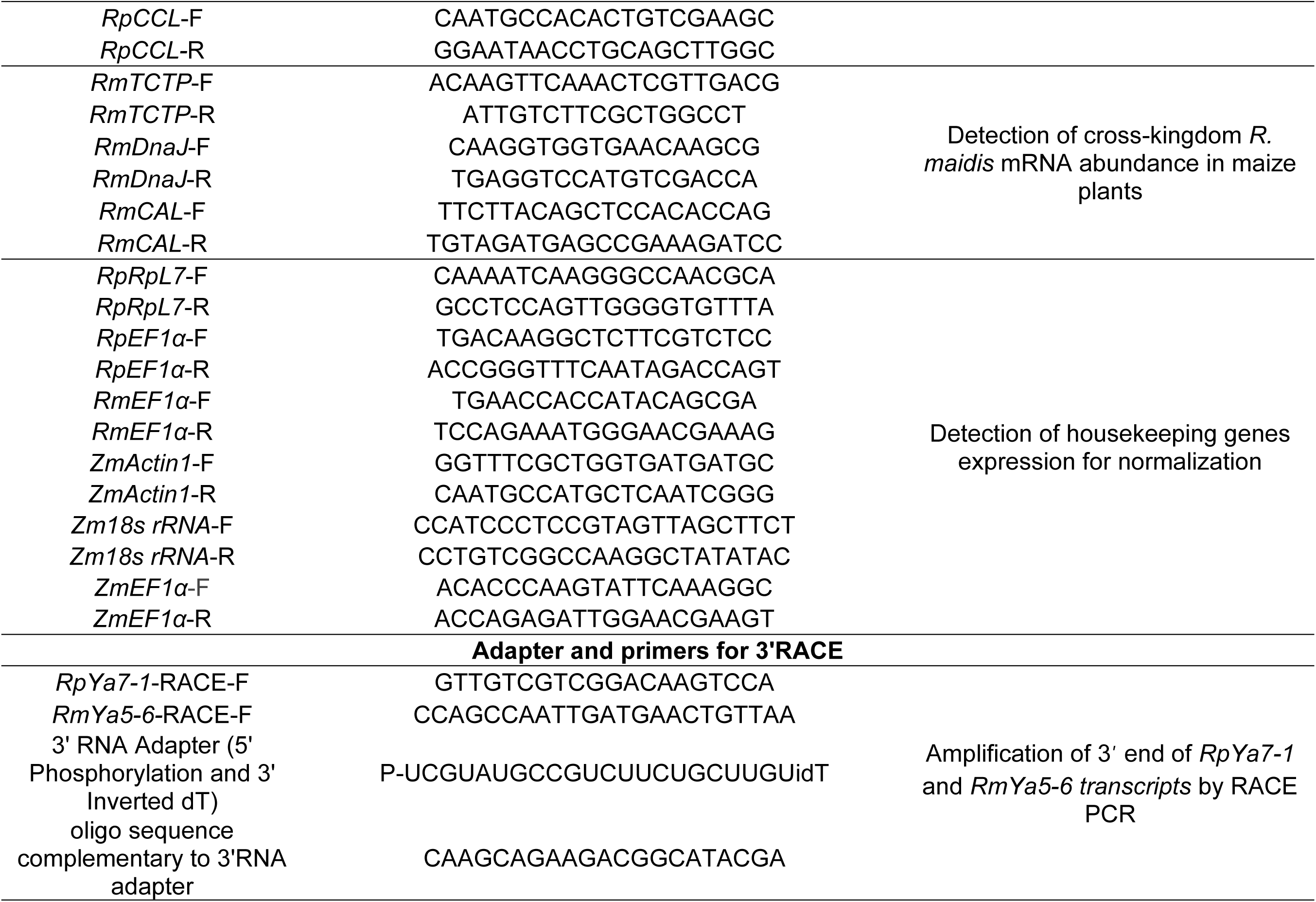
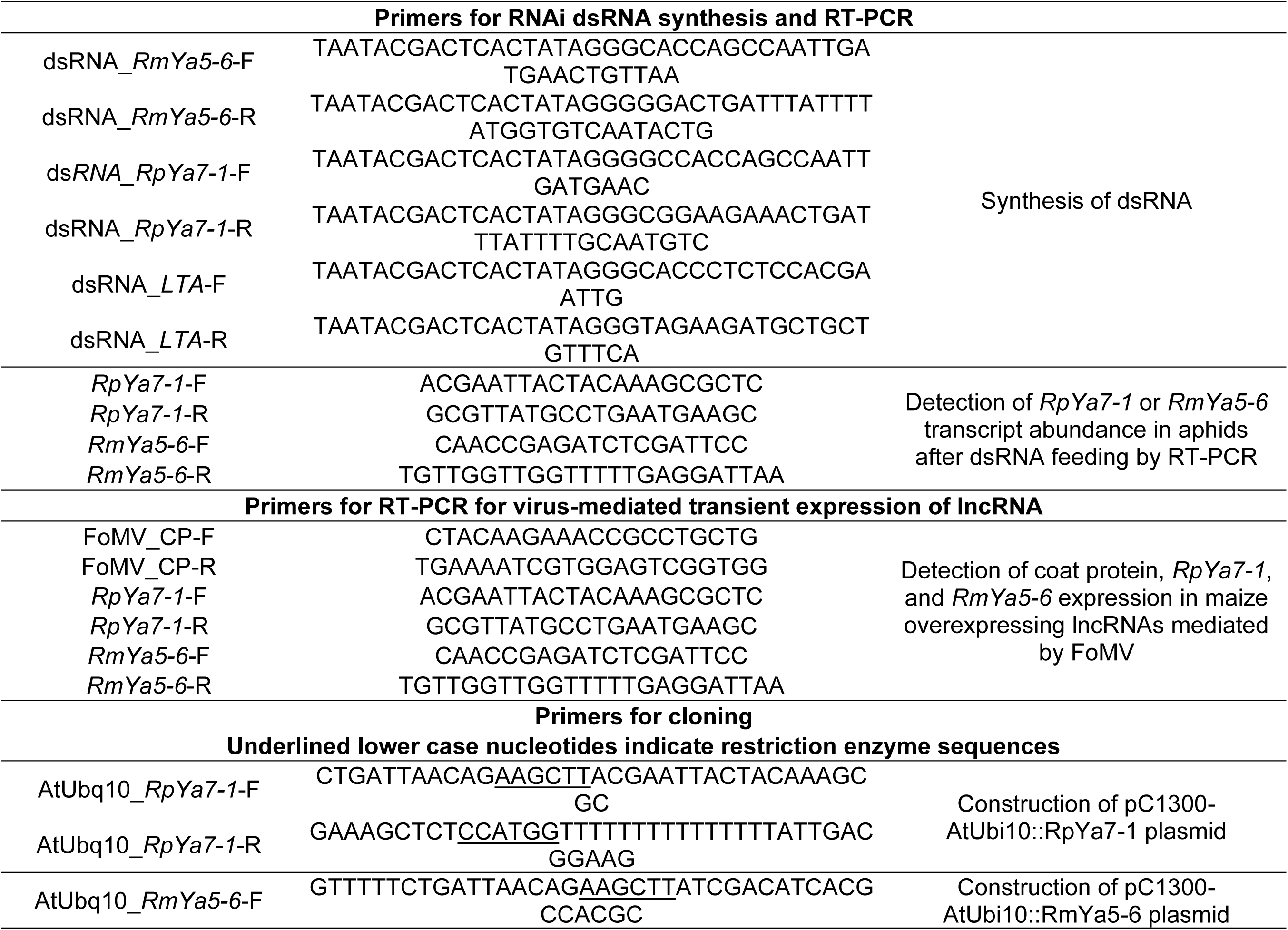
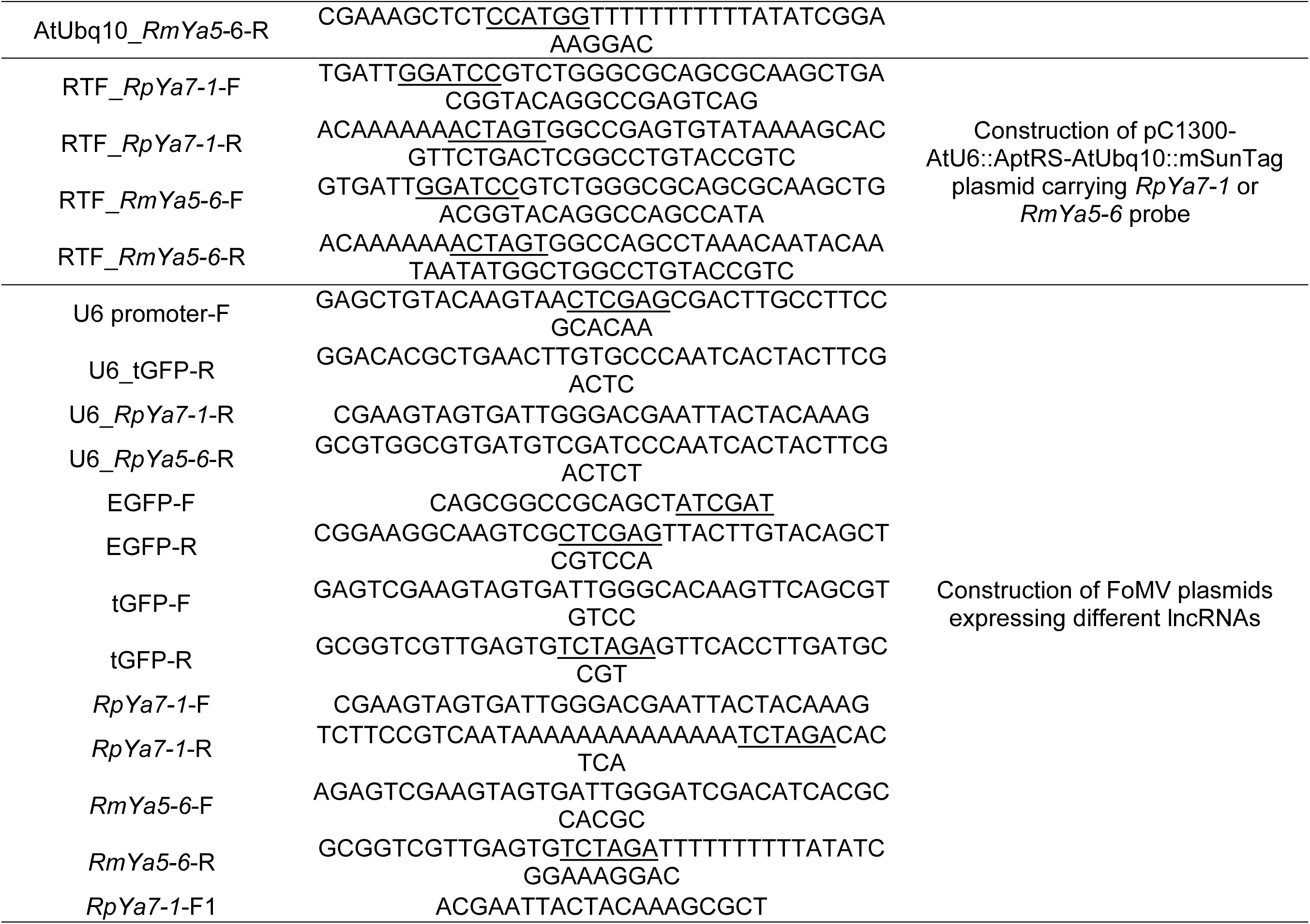
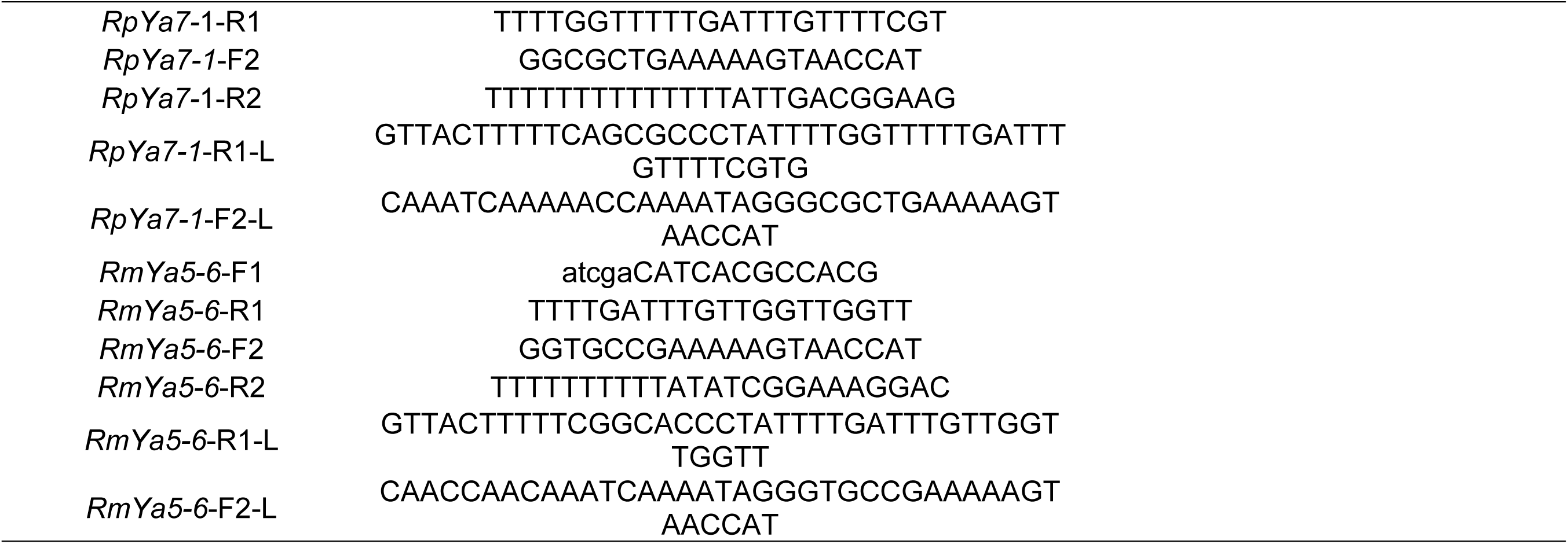
Primers used in this study.

**Table S2.** Coefficient of variation (CV) of abundance in FS for cross-kingdom lncRNAs and mRNAs in *R. padi* and *R. maidis.* This table presents the CV of cross-kingdom lncRNA and mRNA abundance in the FS of *R. padi* and *R. maidis* across 4 replicates. Genes with lower CV values (e.g., <0.5 or 0.8) are considered stably expressed, whereas higher CV values (e.g., >1.0 or 1.2) indicate greater variability in expression. The data correspond to the bar plots shown in Fig. 4C and Fig. 6E.

| Gene | CV | Dataset | Type | Interpretation |
| --- | --- | --- | --- | --- |
| *RpYa1-1* | 0.7209139 | *R.padi* | lncRNA | Moderately stable. Expression is fairly consistent. |
| *RpYa2-1* | 1.6800552 | *R.padi* | lncRNA | High variability. Expression levels differ significantly across the biological samples. |
| *RpYa2-4* | 1.5127367 | *R.padi* | lncRNA | High variability. Expression levels differ significantly across the biological samples. |
| *RpYa3-1* | 0.5185795 | *R.padi* | lncRNA | Stable expression. This gene's expression is relatively consistent across the samples. |
| *RpYa5-3* | 0.9045146 | *R.padi* | lncRNA | Moderate variability. Expression shows some fluctuation between samples. |
| *RpYa6-1* | 1.1340101 | *R.padi* | lncRNA | High variability. Expression levels differ significantly across the biological samples. |
| *RpYa7-1* | 0.7258513 | *R.padi* | lncRNA | Moderately stable. Expression is fairly consistent. |
| *RpYa7-4* | 1.3753409 | *R.padi* | lncRNA | High variability. Expression levels differ significantly across the biological samples. |
| *RpYa7-5* | 0.8904777 | *R.padi* | lncRNA | Moderate variability. Expression shows some fluctuation between samples. |
| *RpES* | 1.1868776 | *R.padi* | mRNA | High variability. Expression levels differ significantly across the biological samples. |
| *RpCDK* | 1.5062946 | *R.padi* | mRNA | High variability. Expression levels differ significantly across the biological samples. |
| *RpCCL* | 1.79067 | *R.padi* | mRNA | High variability. Expression levels differ significantly across the biological samples. |
| *RmYa2-1* | 1.2340522 | *R.maidis* | lncRNA | High variability. Expression levels differ significantly across the biological samples. |
| *RmYa2-3* | 1.1902658 | *R.maidis* | lncRNA | High variability. Expression levels differ significantly across the biological samples. |
| *RmYa3-2* | 1.1620862 | *R.maidis* | lncRNA | High variability. Expression levels differ significantly across the biological samples. |
| *RmYa5-2* | 0.9022982 | *R.maidis* | lncRNA | Moderate variability. Expression shows some fluctuation between samples. |
| *RmYa5-3* | 1.3978512 | *R.maidis* | lncRNA | High variability. Expression levels differ significantly across the biological samples. |
| *RmYa5-4* | 0.8296344 | *R.maidis* | lncRNA | Moderate variability. Expression shows some fluctuation between samples. |
| *RmYa5-5* | 0.8053216 | *R.maidis* | lncRNA | Moderate variability. Expression shows some fluctuation between samples. |
| *RmYa5-6* | 0.4838434 | *R.maidis* | lncRNA | Stable expression. This gene's expression is relatively consistent across the samples. |
| *RmYa6-3* | 0.7983364 | *R.maidis* | lncRNA | Moderate variability. Expression shows some fluctuation between samples. |
| *RmYa6-4* | 0.5759965 | *R.maidis* | lncRNA | Moderately stable. Expression is fairly consistent. |
| *RmYa7-1* | 1.0191401 | *R.maidis* | lncRNA | High variability. Expression levels differ significantly across the biological samples. |
| *RmYa7-2* | 1.2336779 | *R.maidis* | lncRNA | High variability. Expression levels differ significantly across the biological samples. |
| *RmCAL* | 1.765612 | *R.maidis* | mRNA | High variability. Expression levels differ significantly across the biological samples. |
| *RmDnaJ* | 2.3603723 | *R.maidis* | mRNA | Very high variability. Expression is extremely inconsistent between samples. |
| *RmTCTP* | 1.6408967 | *R.maidis* | mRNA | High variability. Expression levels differ significantly across the biological samples. |

**Table S3.** ANOVA Results for Gene × Trait × Time × Plant Part Interactions.

| **Gene** | **Effect** | **NumDF** | **DenDF** | **F-value** | **p-value** |
| --- | --- | --- | --- | --- | --- |
| *RpYa7-1* | Trait:Time | 2 | 229 | 0.7 | 0.496 |
| *RpYa7-1* | Trait:PlantPart | 2 | 229 | 0.61 | 0.542 |
| *RpYa7-1* | Time:PlantPart | 4 | 229 | 2.23 | 0.067 |
| *RpYa7-1* | Trait:Time:PlantPart | 4 | 229 | 0.82 | 0.511 |
| *RpCCL* | Trait:Time | 2 | 229 | 11.64 | <0.001 |
| *RpCCL* | Trait:PlantPart | 2 | 229 | 10.4 | <0.001 |
| *RpCCL* | Time:PlantPart | 4 | 229 | 5.43 | <0.001 |
| *RpCCL* | Trait:Time:PlantPart | 4 | 229 | 8.05 | <0.001 |

**Table S4.** Estimated marginal means (EMMs) and pairwise contrasts of lncRNA and mRNA abundance in maize resistant and susceptible genotypes. Detailed analysis methods are described in *Materials and Methods*. This data corresponds to Fig. 6D.

| Gene | Trait | emmean | SE | Time (h) | Sites |
| --- | --- | --- | --- | --- | --- |
| *RpYa7-1* | Resistant | -6.945154571 | 0.158652768 | 0 | FS |
| *RpYa7-1* | Susceptible | -6.780601964 | 0.250852052 | 0 | FS |
| *RpCCL* | Resistant | -5.359099516 | 0.201339307 | 0 | FS |
| *RpCCL* | Susceptible | -4.288545812 | 0.318345396 | 0 | FS |
| *RpYa7-1* | Resistant | -7.33040108 | 0.158652768 | 0 | UFS |
| *RpYa7-1* | Susceptible | -7.033926618 | 0.250852052 | 0 | UFS |
| *RpCCL* | Resistant | -5.662804525 | 0.201339307 | 0 | UFS |
| *RpCCL* | Susceptible | -5.615579929 | 0.318345396 | 0 | UFS |
| *RpYa7-1* | Resistant | -6.996944702 | 0.158652768 | 0 | DFS |
| *RpYa7-1* | Susceptible | -7.039490993 | 0.250852052 | 0 | DFS |
| *RpCCL* | Resistant | -6.08262132 | 0.201339307 | 0 | DFS |
| *RpCCL* | Susceptible | -5.663232262 | 0.318345396 | 0 | DFS |
| *RpYa7-1* | Resistant | -7.424406467 | 0.158652768 | 12 | FS |
| *RpYa7-1* | Susceptible | -7.041688944 | 0.250852052 | 12 | FS |
| *RpCCL* | Resistant | -5.950731583 | 0.201339307 | 12 | FS |
| *RpCCL* | Susceptible | -6.249548121 | 0.318345396 | 12 | FS |
| *RpYa7-1* | Resistant | -7.048285174 | 0.158652768 | 12 | UFS |
| *RpYa7-1* | Susceptible | -7.295455492 | 0.250852052 | 12 | UFS |
| *RpCCL* | Resistant | -6.259361857 | 0.201339307 | 12 | UFS |
| *RpCCL* | Susceptible | -6.569605938 | 0.318345396 | 12 | UFS |
| *RpYa7-1* | Resistant | -7.240099423 | 0.158652768 | 12 | DFS |
| *RpYa7-1* | Susceptible | -7.252452305 | 0.250852052 | 12 | DFS |
| *RpCCL* | Resistant | -6.70463166 | 0.201339307 | 12 | DFS |
| *RpCCL* | Susceptible | -6.213242942 | 0.318345396 | 12 | DFS |
| *RpYa7-1* | Resistant | -7.392690555 | 0.158652768 | 24 | FS |

| **Gene** | **Trait** | **emmean** | **SE** | **Time (h)** | **Sites** |
| --- | --- | --- | --- | --- | --- |
| *RpYa7-1* | Susceptible | -7.100464782 | 0.250852052 | 24 | FS |
| *RpCCL* | Resistant | -6.537419392 | 0.201339307 | 24 | FS |
| *RpCCL* | Susceptible | -7.004023948 | 0.318345396 | 24 | FS |
| *RpYa7-1* | Resistant | -7.271853207 | 0.158652768 | 24 | UFS |
| *RpYa7-1* | Susceptible | -7.06726646 | 0.250852052 | 24 | UFS |
| *RpCCL* | Resistant | -6.587467377 | 0.201339307 | 24 | UFS |
| *RpCCL* | Susceptible | -6.757839645 | 0.318345396 | 24 | UFS |
| *RpYa7-1* | Resistant | -7.560916847 | 0.158652768 | 24 | DFS |
| *RpYa7-1* | Susceptible | -7.228145571 | 0.250852052 | 24 | DFS |
| *RpCCL* | Resistant | -7.302721739 | 0.201339307 | 24 | DFS |
| *RpCCL* | Susceptible | -6.827359111 | 0.318345396 | 24 | DFS |

## Legends for Datasets S1 to S6

**Dataset S1:** FASTA sequences of *Ya* transcripts identified in *M.persicae*, *R.padi* and *R.maidis*.

**Dataset S2:** GTF annotation file of *Ya* genes from *M.persicae*.

**Dataset S3:** GTF annotation file of *Ya* genes from *R.padi*.

**Dataset S4:** GTF annotation file of *Ya* genes from *R.maidis*.

**Dataset S5:** Transcript TPM values of maize feeding sites in control plants (no aphid) and plants after feeding by *R. padi*.

**Dataset S6:** Transcript TPM values of maize feeding sites in control plants (no aphid) and plants after feeding by *R. maidis*.

## Author contributions

Y.C. and G.W. conceived and designed the study, supervised the project, and finalized the manuscript. J.S. and Z.Z. carried out the majority of the experimental work. C.L., Y.Z., Z.H., and P.Y. contributed to the experimental setup and bioassays. Bioinformatic analyses were conducted jointly by Y.C. and Z.Z. F.W., Y.W., and Y.K. contributed to the performed research. The manuscript was drafted by Y.C., J.S., and Z.Z., with all authors contributing to its revision. All authors reviewed the manuscript, provided feedback, and approved the final version.

## Competing Interest Statement

The authors declare no conflict of interest.

## Data deposition

All RNA-seq data reported in this paper were deposited in the NCBl BioProject under PRJNA1313817. All data are available in the main text or supplementary materials.

## Notes

### Competing Interest Statement

The authors have declared no competing interest.

